# EMCF ecosystem: Towards pretrained foundation model for electron microscopy image analysis

**DOI:** 10.64898/2025.12.09.693109

**Authors:** Zeyu Yu, Jiansheng Guo, Feng Liu, Mengze Du, Shan Xu, Guowei Zhang, Li Xie, Bo Han, Zhonghua Chen, Gaoliang Deng, Chen Rui, Yong He, Xuping Feng

**Affiliations:** College of Biosystems Engineering and Food Science, Zhejiang University, Hangzhou, China; Center of Cryo-Electron Microscopy, Zhejiang University School of Medicine, Hangzhou, China; College of Life Sciences, Biological experiment teaching center, Nanjing Agricultural University, Nanjing, China; Analysis Center of Agrobiology and Environmental Sciences, Zhejiang University.; Department of Computer Science, Hong Kong Baptist University, Hong Kong, China; School of Agriculture, Food & Wine, The University of Adelaide, Adelaide, Australia; Hangzhou Chengfengerlai Digital Technology Co., Ltd, Hangzhou, China; The Rural Development Academy &Agricultural Experiment Station, Zhejiang University, Hangzhou, China

**Keywords:** cellular ultrastructure, foundation model, image segmentation, 3D reconstruction

## Abstract

Volume electron microscopy (vEM) enables nanoscale visualization of three-dimensional (3D) cellular ultrastructure, providing critical insights into physiological processes and pathological alterations. However, its application to large-scale biological tissues remains constrained by two major bottlenecks: prolonged image acquisition and inefficient data processing. Here, we present EMCF ecosystem (EMCFsys), an integrated ecosystem designed to overcome these challenges through three key components: a large-scale benchmark dataset (EMCFD) comprising 4,002,802 high-quality images across 14 EM modalities and 6 biological kingdoms; a foundation image restoration model (EMCellFiner); and a scalable image analysis foundation model (EMCellFound). Together, these modules systematically enhance image quality and substantially improve analysis efficiency. Our results show that EMCellFiner outperforms specialist models in restoring degraded images, even surpassing original ground truth sharpness in certain artifact regions, and reduces imaging time by 16-fold by enabling low-resolution and low dwell time acquisition. EMCellFound exhibits exceptional feature discriminability, outperforms specialist models in classification, semantic segmentation and instance segmentation. It also enables high-precision 3D reconstruction of organelles (e.g., endoplasmic reticulum) with minimal labeled data (0.01% of total volume). We validated the EMCFsys on unseen datasets across diverse biological contexts and imaging platforms. By publicly releasing both the dataset and models, we establish a scalable paradigm for automated, high-throughput vEM data interpretation, accelerating exploration of life’s nanoscale structure and function across biology.

## Main

Compared with cultured cells, in situ nanoscale three-dimensional (3D) ultrastructural analysis of cells, tissues, and even organs within biological specimens are essential for a profound understanding of biological function, particularly for elucidating mechanisms underlying clinical diagnosis and pathological detection^1,2^. For instance, the three-dimensional architecture of the endoplasmic reticulum (ER) is highly dynamic, with its tubular and sheet-like domains undergoing adaptive remodeling that is intimately linked to cellular homeostasis and disease progression^3^. Notably, ER fragmentation, observed in neurodegenerative disorders, represents a critical driver of neuronal death^4^.

As a key technology driving the next revolution in biology^5^, vEM has already enabled breakthrough discoveries across numerous biological frontiers^6–8^. currently, this technique stands out as a pioneering approach uniquely capable of achieving computational reconstruction of nanoscale ultrastructure across substantial biological volumes, including whole brains^9,10^. The global vEM research community is striving to leverage this technology to visualize, at nanometer resolution, the complex ultrastructure of organelles, cells, and tissues at the micrometer scale, thereby elucidating their intricate interactions with the surrounding micro-environment and enabling the construction of large-scale 3D atlases of biological specimens. However, imaging large tissue volumes remains extremely time-consuming. For example, a complete mouse brain volume would require an estimated 3.6 million years per microscope under conventional acquisition settings^11^. Furthermore, the lack of efficient, automated image processing pipelines continues to pose a major bottleneck to the broad application of vEM in life sciences.

Among these limitations, image processing has emerged as an even greater bottleneck than image acquisition. The challenges primarily arise from two sources: first, the intrinsic properties of electron microscopy (EM) data, such as high dimensionality, complex textures, noise, low contrast, and imaging artifacts, severely impede effective information extraction. Second, biological structures themselves exhibit immense variability; the morphology and scale of organelles differ significantly across species, cell types, and even within the same cell under different physiological states, making it exceedingly difficult to develop generalizable recognition models.

Although deep learning-based automated analysis methods have made substantial advances in specific tasks, such as image enhancement, super-resolution, organelle segmentation^2,12–15^, current approaches are nevertheless constrained by fundamental limitations. Mainstream supervised learning models are highly dependent on large-scale, high-quality manual annotations, a process that is not only time-consuming and laborious but also yields models that are often task-specific and generalize poorly to new species or biological contexts^16^. Researchers are therefore frequently forced into tedious trade-offs and combinations of manual, semi-automated, and fully automated methods, drastically constraining the efficiency of vEM-driven discovery.

To overcome this critical bottleneck, we draw inspiration from “Foundation Models”, which have achieved remarkable success in computer vision. By pretraining on massive and diverse datasets, foundation models learn general-purpose, transferable feature representations that exhibit exceptional performance and generalization across a wide range of downstream tasks^17–19^. However, the primary obstacle to applying this powerful paradigm to cellular EM is the current lack of a unified, large-scale, diverse, and high-quality pretraining dataset.

To address these limitations, we introduce EMCFsys that comprises three core components: (1) the EMCF dataset (EMCFD), a large-scale cellular EM benchmark we curated, encompassing multiple species, cell types, and imaging modalities; (2) EMCellFiner, an image restoration and enhancement foundation model pretrained on EMCF dataset, that also used for reduces imaging time; and (3) EMCellFound, a powerful and generalizable segmentation foundation model capable of precisely delineating a multitude of organelles and subcellular structures. Our results show that this data-driven foundation model approach not only systematically improves vEM image quality but also enables unprecedented zero-shot or few-shot segmentation across species and cell types. The EMCFsys provides a scalable, new paradigm for the high-throughput, automated interpretation of vEM data, paving the way for exploring the mysteries of life’s structure and function on a vastly broader biological scale.

## Results

### Overview of EMCF ecosystem

The EMCFsys has three components (Fig. 1): (i) the Electron Microscopy Cell Foundation Dataset (EMCFD), a vast and diverse repository of curated 4,002,802 cell images; (ii) EMCellFiner, a foundation model engineered for high-fidelity image restoration; and (iii) EMCellFound, a versatile foundation model designed for seamless adaptation to downstream analytical tasks. EMCellFiner restores low-resolution or noisy EM images to exceptional clarity, while EMCellFound facilitates robust transfer learning for applications such as image classification, semantic and instance segmentation, and rapid 3D reconstruction. To rigorously validate the generalizability of our models, we benchmarked their performance on multiple unseen datasets, sourced from a variety of biological contexts and imaging platforms. The EMCFsys is now publicly accessible to facilitate broader researcher engagement with our models, data and code. To enhance usability within the Amira platform, we have developed plugins based on EMCellFiner and EMCellFound.

**Fig. 1.**
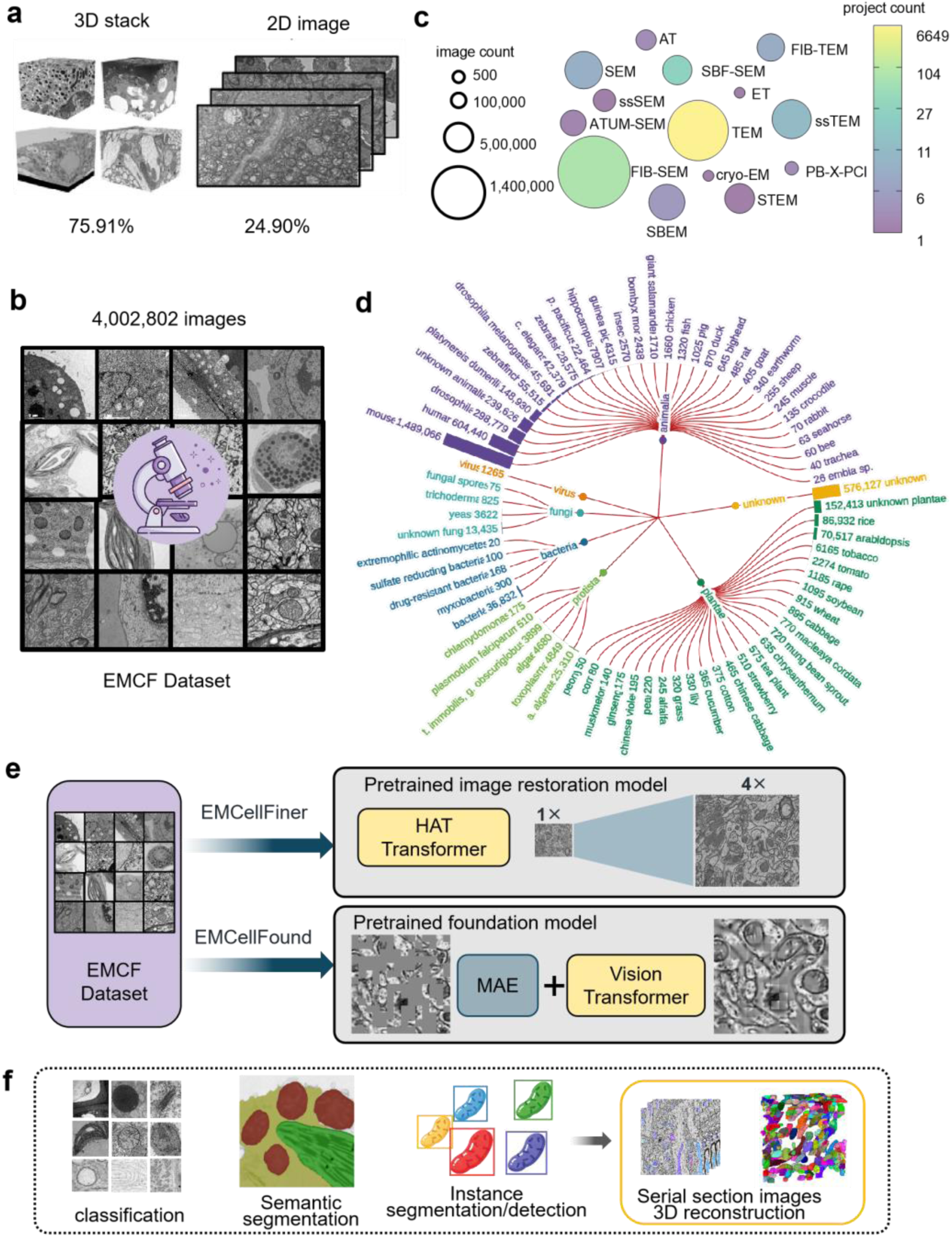
Overview of Electron Microscopy Cell Foundation (EMCF). **a**, The raw data includes both 3D and 2D electron microscopy images, offering a comprehensive representation of cellular structures. **b,** The EMCF dataset, a large-scale and diverse pretraining dataset comprising 4,002,802 electron microscopy images. **c,** Distribution of images across 14 electron microscopy acquisition methods and their respective data volume proportions. **d,** Taxonomic distribution of the specimens from which electron microscopy images were acquired, covering a broad spectrum of biological species. **e,** Overview of the foundational models: EMCellFiner, an image restoration foundation model, and EMCellFound, a general-purpose foundational large model designed for adaptation to various downstream EM analysis tasks. **f,** Adaptation of the EMCellFound foundational model to diverse downstream tasks, including organelle classification, semantic segmentation, instance segmentation, and 3D reconstruction.

### A curated foundation dataset for cellular electron microscopy

Our EMCFD is the first large-scale, multi-domain foundation dataset specifically designed for cellular EM. Its raw data was aggregated from two primary channels: millions of images acquired in our laboratory and a vast collection of 2D and 3D images compiled from 6,825 public repositories, studies, and publications (Fig. 1a-d). Recognizing the inherent heterogeneity in magnification, resolution, and quality across these sources, we implemented a systematic curation pipeline to filter, clean, and standardize the data (Supplementary Fig. 1). This process yielded the final EMCFD, comprising 4,002,802 high-quality electron micrographs. The dataset exhibits substantial diversity across 14 distinct electron microscopy modalities, including transmission electron microscopy (TEM), scanning electron microscopy (SEM), focused ion beam scanning electron microscopy (FIB-SEM), and cryo-electron tomography (Cryo-ET), among others (Fig. 1c). Biologically, it encompasses six major kingdoms: Animalia, Plantae, Fungi, Protista, Bacteria, and Viruses (Fig. 1d). Images derived from Animalia constitute the largest proportion (81.29%), predominantly from key model organisms such as mouse, human, and Drosophila. Plantae-derived images account for 8.90%, featuring model species and crops including Arabidopsis thaliana, rice, and tobacco. The remaining dataset fraction includes images from Bacteria, Fungi, and Viruses, as well as a subset of samples classified as “unknown.” This unprecedented data resource, with its high quality, broad species representation, and multi-modal technical scope, provides a solid foundation for developing the next generation of AI models in cell biology.

### EMCellFiner for super-resolution

EMCellFiner, a foundation model engineered for 4× super-resolution, achieves high-fidelity restoration of degraded electron microscopy (EM) images by leveraging a HAT Transformer architecture trained on the EMCF dataset (Fig. 2a). To systematically evaluate our EMCellFiner foundation model, we first assessed its performance in a supervised setting where ground truth images were available. We simulated common image degradation by applying a 5×5 Gaussian blur and 4-fold downsampling to high-quality source images. In comparisons, EMCellFiner consistently outperformed the specialist model EMDiffuse^12^, generating visually sharper reconstructions (Fig. 2c) and achieving superior quantitative scores across both Peak Signal-to-Noise Ratio (PSNR) and Feature Similarity Index^20^ (FSIM) metrics (Fig. 2b). Moreover, we demonstrated that EMCellFiner exhibits robust restoration capabilities across a wide range of degradation types, including various blurs and noises (Fig. 2d and Supplementary Figs. 2-3). Remarkably, its reconstructions often surpassed the sharpness of the original ground truth images. For instance, after 3×3 Gaussian blur degradation, EMCellFiner’s output was 1.29-fold sharper than the original ground truth images, whereas EMDiffuse’s output was only 0.57-fold (Supplementary Fig. 2a). This superior performance extended across various challenging scenarios (Supplementary Figs. 4-5) and organelles (Supplementary Figs. 6-8). For instance, EMCellFiner correctly reconstructed the nuclear envelope and chromatin structure where EMDiffuse introduced distortions. Furthermore, it demonstrated immunity to strong image artifacts, such as bright stripes in ER images, that caused failures in the EMDiffuse reconstruction. These results indicate that EMCellFiner possesses greater stability and reliability across different cellular contexts.

**Fig 2.**
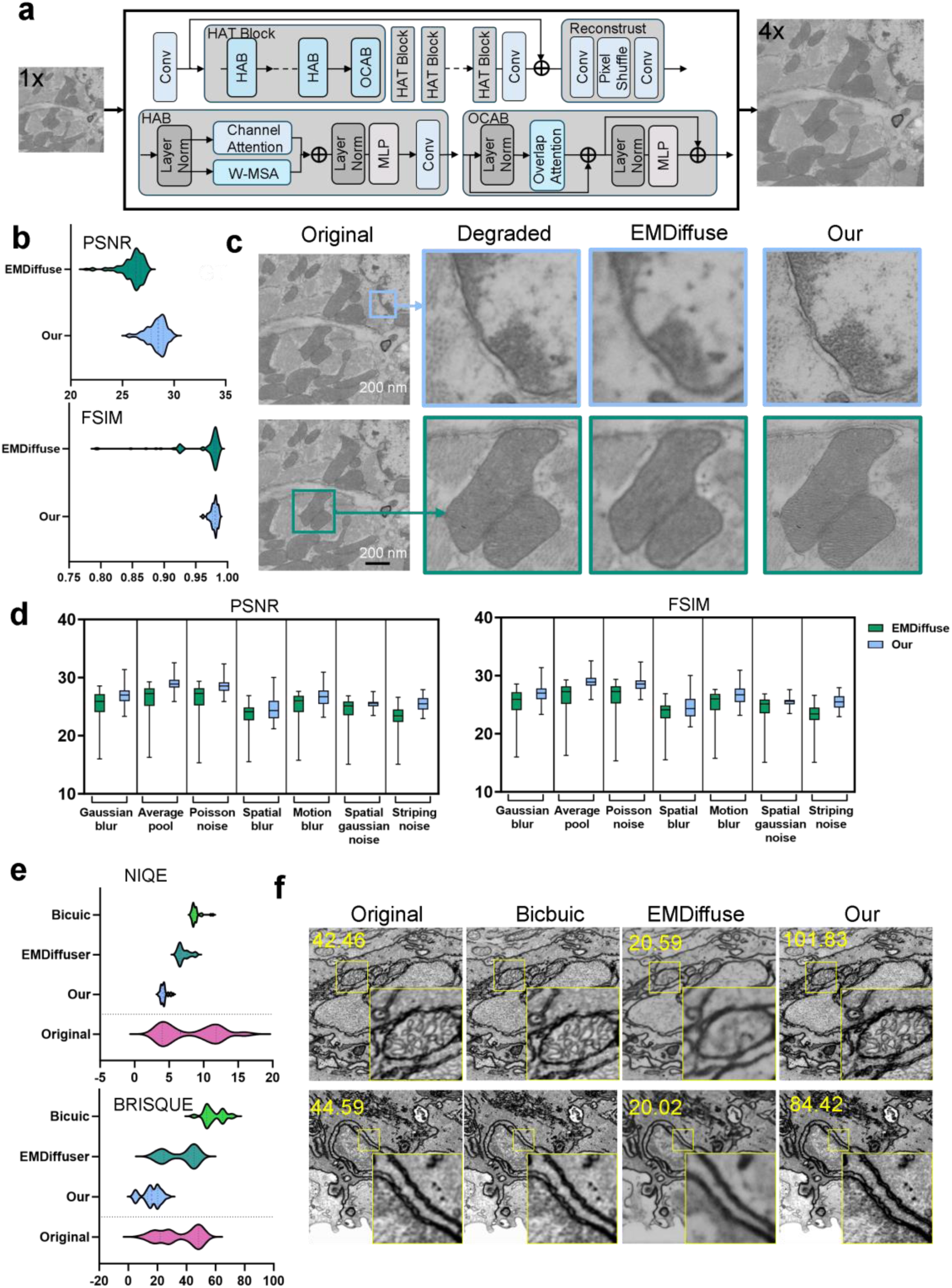
EMCellFiner method and super-resolution performance evaluation. **a,** The architecture of EMCellFiner. Training input samples are synthetically degraded using 4x downsampling and Gaussian blur. **b,** Comparison of PSNR and FSIM for super-resolution (SR) reconstructions generated by EMCellFiner and EMDiffuse under high Signal-to-Noise Ratio conditions, specifically for images degraded by Gaussian blur. **c,** Representative SR images reconstructed by EMCellFiner and EMDiffuse. Input images for these examples were degraded by a 5×5 Gaussian blur. **d,** Comparison of PSNR and FSIM for SR reconstructions generated by EMCellFiner and EMDiffuse across various degradation patterns. **e,** Comparison of BRISQUE and NIQE for SR reconstructions by Bicubic interpolation, EMCellFiner, and EMDiffuse under low Signal-to-Noise Ratio conditions. **f,** Representative SR images reconstructed by EMCellFiner and EMDiffuse under low Signal-to-Noise Ratio conditions. Sharpness values are shown for related images. The data sources used for these tests are detailed in Supplementary Table 1.

To assess real-world applicability where ground truth images are often unavailable, we evaluated EMCellFiner’s zero-shot generalization capability on diverse unseen datasets. Using the no-reference quality metrics BRISQUE and NIQE as valuation indexes, EMCellFiner significantly outperformed both standard bicubic interpolation and EMDiffuse (Fig. 2e). Visually, the model enhanced clarity and restored critical biological features, such as organelle membranes and cell junctions, without introducing new artifacts (Fig. 2f). This robust performance was consistent across datasets originating from various EM instruments, further confirming its wide-ranging utility (Supplementary Fig. 9).

Collectively, these results establish EMCellFiner as a powerful, generalist foundation model for EM image restoration. Its ability to restore high-fidelity images from lower-quality inputs has profound practical implications for experimental design. By enabling 4× super-resolution, researchers can acquire images at a lower resolution (e.g., 512×512 instead of 2048×2048), reducing acquisition time by a factor of approximately 16-fold, for example, from 16 hours to just 1 hour, without compromising the final image quality. This acceleration dramatically increases experimental throughput and reduces operational costs.

### EMCellFound learns discriminative features for organelle classification

We developed EMCellFound, a generalist foundation model for cellular feature representation, by employing a Masked Autoencoder^21^ (MAE) self-supervised strategy on EMCFD (Fig. 3a). To benchmark its performance, we compared MAE against leading self-supervised learning methods (DINO^22^, MoCov3^23^, and All4One^24^) on a challenging 8-class organelle classification task. With the pretrained Vision Transformer backbone frozen, a k-nearest neighbors^25^ (KNN) classifier was applied to the learned features. Under identical training conditions, the MAE-pretrained EMCellFound achieved a mean accuracy of 96.01%, significantly outperforming all other strategies (Fig. 3b). To qualitatively assess the learned representations, we visualized the feature space using t-SNE^26^ (Fig. 3c). The visualization revealed well-defined and distinct clusters corresponding to the eight organelles classes, corroborating the model’s robust discriminative capacity. Notably, we observed a partial overlap between the clusters for the Golgi and endosomal. This finding is consistent with their known biological relationship, as both organelles share similar vesicular morphologies and are functionally linked through intracellular trafficking pathways^27^.

**Fig. 3.**
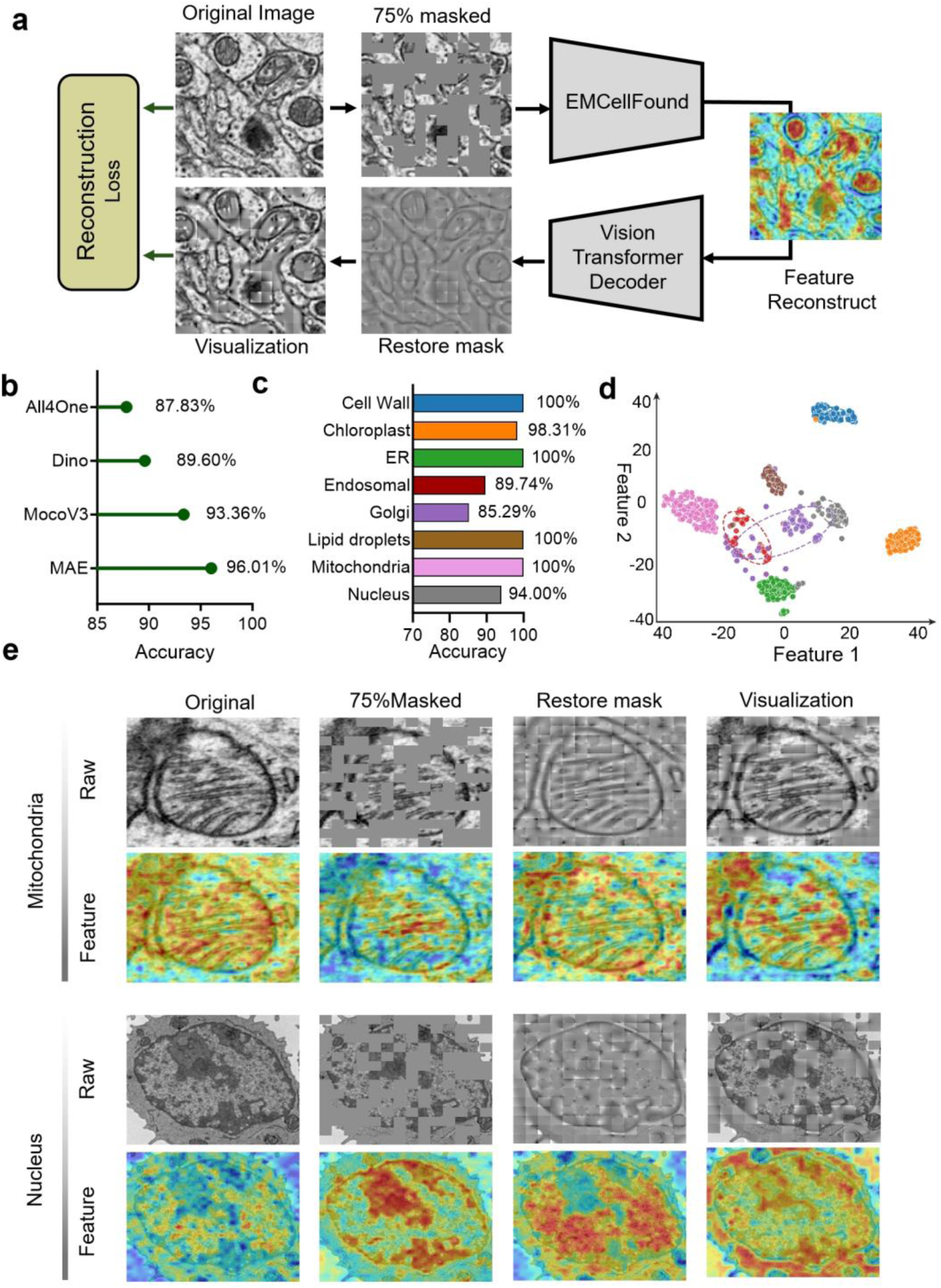
EMCellFound Performance in Zero-Shot Organelle Classification. **a**, Architecture of the EMCellFound model based on the Masked Autoencoder (MAE) self-supervised learning framework. **b**, Zero-shot KNN classification accuracy comparison of ViT base models pretrained with different self-supervised algorithms (MAE, DINO, MoCo v3, All4One). **c**, t-SNE visualization of feature distributions for eight organelle classes, extracted from the EMCellFound encoder. Endoplasmic reticulum (ER). **d**, Classification performance (precision) across the eight organelle classes using EMCellFound. In panels **c** and **d**, organelle categories are distinguished using the same color scheme. **e**, Feature visualization for mitochondria and nucleus recognition using Class Activation Mapping. Upper row: I Input image processing pipeline (raw image → 75% masked input → reconstructed feature map → final reconstruction). Bottom row: Attention heatmaps from Transformer layers 3, 6, 9, and 12, highlighting maximal attention regions. Color bars were generated from the importance scores of a specific prediction learning by the modules, displayed in different colors from blue to red, indicating increasing importance scores.

The efficacy of the MAE approach was evident in the model’s ability to reconstruct fine ultrastructural details from inputs where 75% of image patches were randomly masked (Fig. 3e). To probe the model’s internal representations, we visualized feature activation maps from successive layers of the Transformer architecture (Supplementary Figs.11-12). This analysis revealed a distinct learning hierarchy: shallow layers were sensitive to generic, low-level features such as membrane contours, while deeper layers focused on higher-level, semantically meaningful regions of high electron density.

Crucially, EMCellFound learned highly discriminative and biologically relevant features for each organelle class. For instance, its attention mechanism correctly highlighted the nuclear envelope, internal chromatin, and the nucleolus when analyzing nuclei. Its focus was similarly specific for other organelles, targeting the folded cristae of mitochondria, starch granules within chloroplasts, and the unique vesicular patterns of endosomes (Supplementary Figs. 11, 12). This remarkable correspondence between the model’s learned features and the morphological criteria used by expert cell biologists underscores the power of EMCellFound to capture meaningful biological information.

### EMCellFound can transfer to semantic segmentation

To assess the transferability of EMCellFound to downstream tasks, we benchmarked its performance on the Plantorganelle Hunter dataset, covering four distinct organelle types, covering four distinct organelle types. We compared various fine-tuning strategies (Supplementary Figs. 13) against scratch-trained baselines (Unet^28^, ViT^29^, Mask2former^30^) and the domain-specialist model, OrgSegNet^15^.

Leveraging the pretrained representations, EMCellFound model consistently outperformed all baselines. Under full fine-tuning, it achieved the highest mIoU, surpassing the specialist OrgSegNet (Fig. 4a and Supplementary Figs. 14) while demonstrating significantly faster convergence, requiring fewer training epochs to reach optimal performance (Fig.4b).

**Fig 4.**
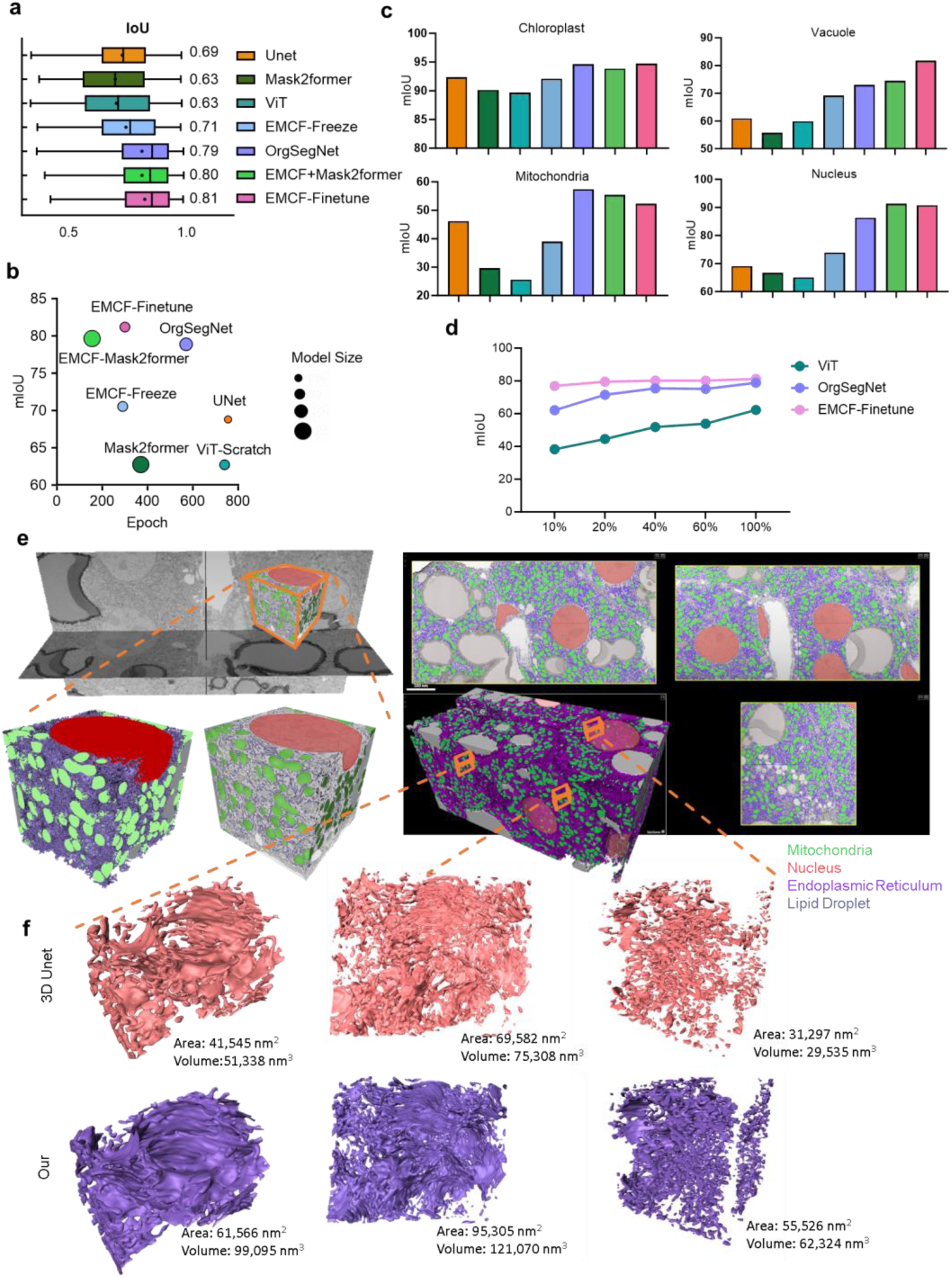
EMCellFound performance in downstream semantic segmentation task. **a**, Semantic segmentation results of four baseline models (Unet, ViT, Mask2Former, OrgSegNet) and three fine-tuned EMCF (EMCF-Freeze, EMCF-Mask2Former, and EMCF-Finetune) on the Plantorganelle Hunter dataset. **b**, Comparison of segmentation performance across seven models based on the mIoU, Epoch and model size in the inference process. **c**, Organelle-specific segmentation performance across seven models. **d**, Segmentation results of ViT, OrgSegNet, and EMCF-Finetune with increasing training dataset size. e, Representative 3D reconstruction of organelles from Liver-6 vEM data using EMCF-Finetune. A selected volumetric regionfrom the Liver-6 vEM dataset is segmented in 2D slices by EMCF-Finetune and subsequently reconstructed into a 3D model. Lipid droplets (LD), endoplasmic reticulum (ER), mitochondria, and nuclei are shown in gray, purple, green, and translucent red, respectively. f, Comparative 3D reconstruction of the intricate ER network from e using a baseline 3D U-Net (top, red) and EMCF-Finetune (bottom, purple). Quantitative metrics (total surface area and volume) highlight the improved topological completeness and smoothness achieved by EMCF-Finetune, overcoming fragmentation issues seen in the baseline. Note: **a,** Unet, Mask2former(ViT as backbone with Mask2former as segmentation head), ViT (ViT as backbone with UperNet^32^ as segmentation head) are stratch-trained models. EMCF-Freeze is the model with the backbone weight freezing, and only train the UperNet segmentation head. EMCF-MaskFormer is the model consist of EMCellFound backbone with Mask2former segmentation head, and EMCF-Finetune is the model with EMCellFound backbone with Uperhead segmentation head. see Supplementary Fig. 13.

We compared the segmentation performance of the models across different organelles (Fig.4c). Overall, the EMCellFound weight injection model (EMCF-Finetune and EMCF-Mask2former) demonstrated comparable performance to the specialized model across various organelles. For chloroplasts, which are abundant in the dataset, all models achieved an IoU exceeding 90%. For the class-imbalanced and less numerous Nucleus, “full fine-tuning” surpassed the specialized model. Additionally, EMCF-Finetune achieved peak performance with only 10% of training data (Fig. 4d), outperforming both scratch-trained ViT (requires 100% data) and OrgSegNet (requires 60% data) under equivalent conditions. This confirms that the foundation model effectively transfers learned structural priors, drastically reducing the annotation burden for downstream tasks.

We further extended EMCellFound to the challenging task of 3D reconstruction from sparse 2D annotation slices, which requires fewer GPU resources compared to directly segmenting organelles in 3D vEM data. Using the Liver-6 vEM dataset^31^, we evaluated whether our model could reconstruct complex organelle structures with minimal computational and annotation costs. Strikingly, trained on only 0.01% of the total volume, EMCellFound reconstructed high-fidelity 3D models of mitochondria, nuclei, and lipid droplets, significantly outperforming the standard 3D U-Net (Fig. 4e,f). The advantage was most pronounced for the ER, which is the smallest target within the entire field of view, characterized by indistinct features easily confused with others. While 3D U-Net struggled with fragmentation, EMCellFound produced topologically continuous, smooth, and complete surfaces (Supplementary Fig. 15), preserving the integrity of this intricate membrane network. Both their volume and surface area metrics showed improvements compared to 3D-Unet (Fig. 4f).

### Foundation pretraining enables few-shot instance segmentation

To evaluate the generalization capability and data efficiency of our proposed model for instance segmentation, we conducted benchmark tests on three public electron microscopy datasets of mitochondria: Bock^33^, Kidney^34^, and Lee^35^ (Fig. 5). We systematically compared models initialized with EMCellFound weights against equivalent architectures trained from scratch, encompassing both standard CNN backbone (RTM^36^) and Transformer-based (ViT and ViTAdapt^37^) backbones (several model configurations can be seen in Supplementary Fig. 16). The results revealed that the standard RTM model, when trained from scratch, substantially underperformed all other models in both segmentation and detection tasks. Critically, the scratch-trained ViT-RTM and ViTAdapt-RTM models also consistently yielded lower mean Average Precision (mAP) for instance segmentation across all three datasets compared to our foundation models initialized with pretrained weights (Fig. 5a), a finding corroborated by the visual results in Fig. 5c. The integration of pretrained weights in EMCF-Finetune and EMCFAdapt-Finetune led to a significant mAP increase for detecting both small and large mitochondria (Fig. 5b).

**Fig. 5.**
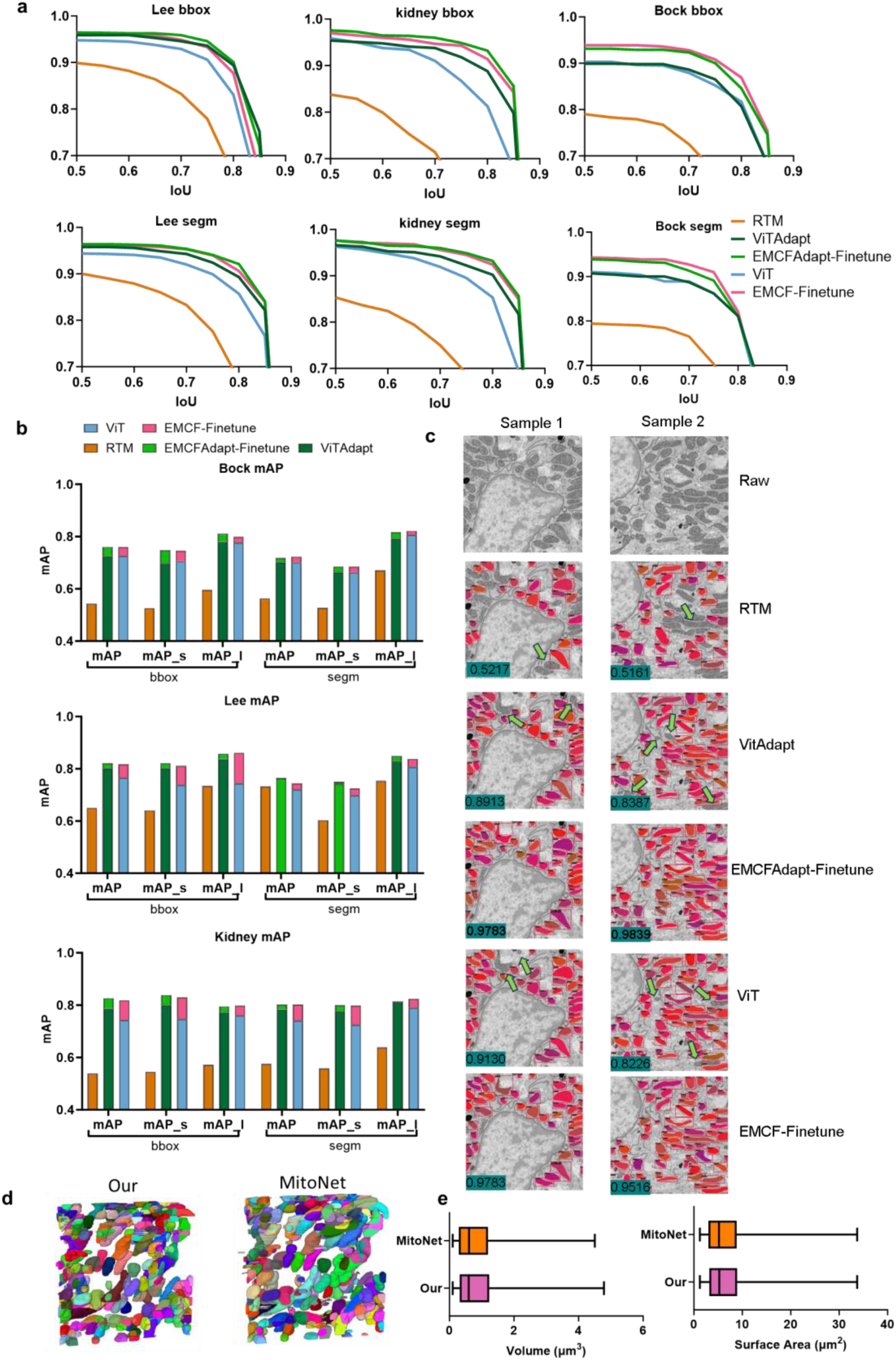
Application of EMCellFound for instance segmentation of mitochondria. **a,** Mitochondrial instance segmentation results of five models on Bock, Lee, and kidney datasets. The first and second columns represent the mean Average Precision (mAP) for bounding box detection and segmentation, respectively. The strategies of the five models are detailed in Supplementary Fig. 17. **b,** Detection and instance segmentation performance of models across different object sizes. mAP, mAP, and mAP denote the mean Average Precision, mAP for small objects (bounding box size < 96*96 pixels), and mAP for large objects (bounding box size > 96*96 pixels), respectively. c, Representative instance segmentation of mitochondria by five models on kidney data. The numbers within the images represent the bounding box recall (Recall = correctly detected instances / total ground truth instances). **d,** Results of 3D Mitochondrial reconstruction from Kidney data following aumomatic segmentation by EMCF and MitoNet. **e,** Relations of surface area and volume of mitochondria obtained from EMCF and MitoNet. (n = xxx independent mitochondrias, mean ± s.d.).

Furthermore, we benchmarked our model against MitoNet^14^, a state-of-the-art, specialized model for mitochondrial instance segmentation trained on an extensive dataset of 21,860 images. Remarkably, by fine-tuning our EMCellFound model on only 30 images (1024×1024 pixels) from the Kidney dataset, representing a mere 0.14% of the data used to train MitoNet, we achieved performance parity with this heavily trained specialist model (Figs. 5d,e). These findings underscore a transformative shift in paradigm that EMCellFound eliminates the need for prohibitive large-scale dataset curation, delivering high-precision analysis with minimal supervision.

### Enables Efficient 3D reconstruction of human anterior pituitary cells

High-resolution volumetric imaging of large biological specimens involves a prohibitive trade-off between acquisition speed and structural fidelity. While vEM techniques like FIB-SEM offers nanoscale precision, acquiring dense volumes is notoriously time-consuming. EMCellFiner breaks this trade-off. By computationally upscaling the rapid low-resolution scan, it reconstructs a high-fidelity 4 times upsacle volume in 6 hours of offline computation using a RTX3090 GPU (Fig. 6b). This workflow liberates valuable microscope resources, effectively reducing instrument time and associated costs by 94% (16-fold). For instance, imaging a stack of 854 high-resolution images (4096×3536 pixels) of anterior pituitary cells, with a pixel dwell time of 5 μs, takes approximately 32 hours. In contrast, under identical settings, acquiring a stack of 854 images at lower resolution (1024×884 pixels) requires only 2 hours. Compared to the EMDiffuse model, EMCellFiner demonstrates superior performance on unseen anterior pituitary data, achieving faster reconstruction times and higher quality outputs (Fig. 6c-f). The enhanced sharpness and clarity of organelle membranes produced by EMCellFiner facilitate more accurate downstream analysis and segmentation.

**Fig. 6.**
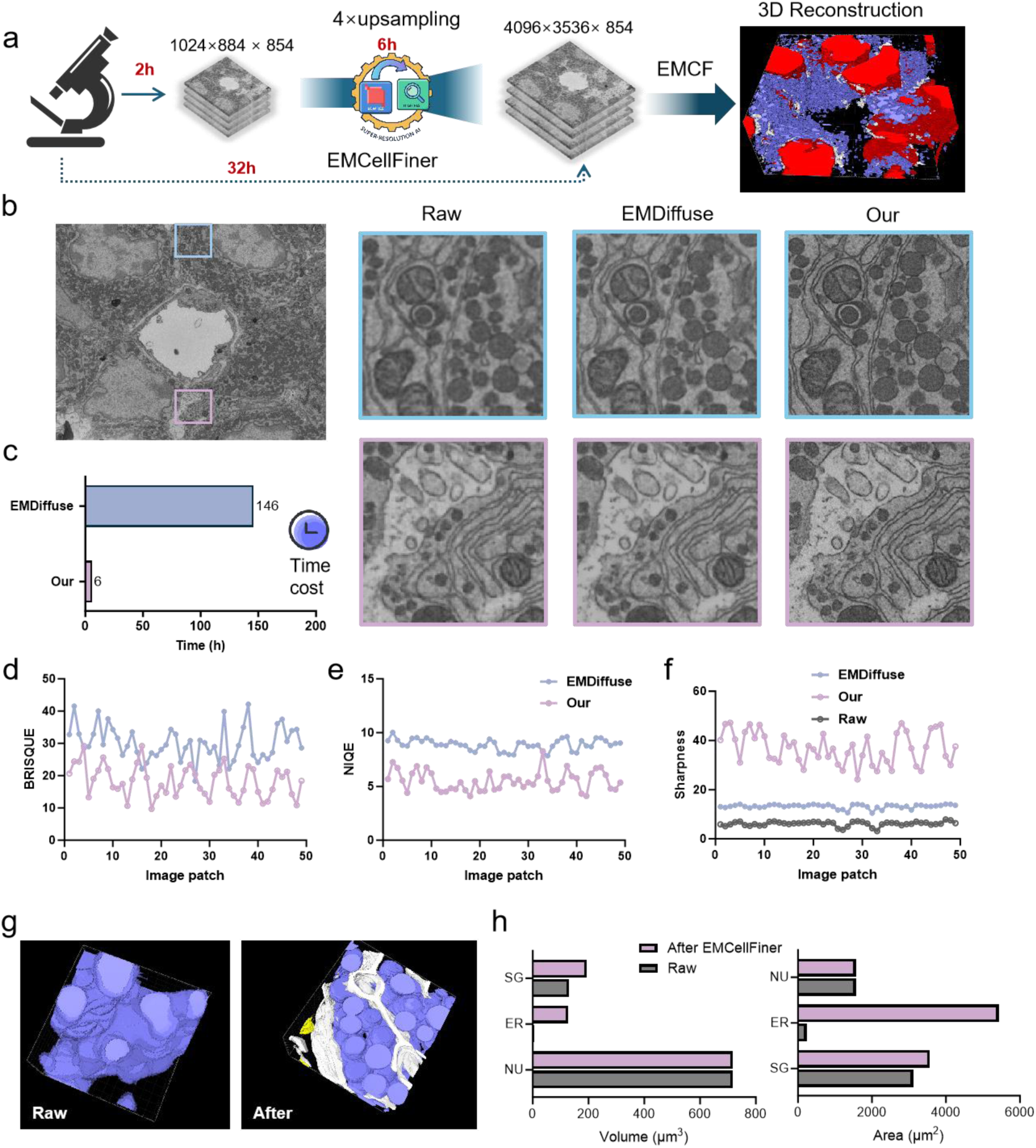
Performance of EMCellFiner and EMCF pipline for 3D reconstruction of human anterior pituitary cells. a, Workflow schematic showing reduced total acquisition time by combining rapid low-resolution (LR) imaging with EMCellFiner’s super-resolution (SR) capability. **b,** zoomed-in representative SR images of anterior pituitary cells processed by EMCellFiner and EMDiffuse. **c,** Comparison of the computational time required by EMCellFiner and EMDiffuse to upscale image stacks from 1K to 4K resolution. On an NVIDIA RTX 3090 GPU, EMCellFiner completes high-fidelity super-resolution reconstruction in just 6 hours (approximately 0.5 frames per minute at 1k resolution), whereas EMDiffuse requires about 146 hours (approximately 0.02 frames per minute) under the same conditions. **d-f**, Comparison of image quality metrics for patches generated from a large-scale image using EMCellFiner and EMDiffuse. The evaluation uses (**d**) NIQE, (**e**) BRISQUE, and (**f**) sharpness scores. **g,** Comparison of 3D cell models (4096×3536×854) reconstructed from original LR data and EMCellFiner-enhanced SR data. Purple is secretory granules, white is endoplasmic reticulum. **h,** Quantitative comparison of surface area and volume for the nucleus (NU), endoplasmic reticulum (ER), and secretory granules (SG), derived from LR- and SR-based reconstructions.

To demonstrate the synergy between restoration model EMCellFiner and foundation model EMCellFound, we employed a few-shot learning strategy, fine-tuning EMCellFound on just one randomly selected slice to segment the entire 854-slice volume. EMCellFiner significantly improved segmentation accuracy, particularly for the ER. In the 1K raw images, ER structures were undetectable; however, EMCellFiner processing enabled the EMCellFound model to reliably segment these intricate networks within the 2D stack images. Visualization in Arivis 3D revealed clear connections and spatial distributions between the ER, secretory granules, mitochondria, and the nucleus of anterior pituitary cell (Supplementary Fig. 17 and 18). This result reveals a critical distinction that while prominent structures, like nuclei and secretory granules, are robust to low resolution, restoration to high quality is indispensable for capturing fine-grained, irregular organelles such as the ER. Consequently, processing with EMCellFiner yielded a substantial quantitative recovery of biological signal, reflected in the marked increase in detected organelle volume and surface area (Fig. 6h).

## Discussion

In this work, we introduce EMCellfiner and EMCellfound, two self-supervised foundation models specifically designed for EM image analysis. Complementing these models is the EMCF dataset, a large-scale, multi-species, and multi-modal cellular EM image collection comprising 4,002,802 images. Together, this integrated ecosystem provides a comprehensive solution to address critical challenges in the end-to-end analysis of EM data. Our findings demonstrate that the synergistic integration of EMCellfiner and EMCellfound significantly reduces imaging time, requiring only minimal fine-tuning on a few images to achieve accurate organelle segmentation for large cell tissue (Fig. 6).

Large scale tissue imaging remains extremely time-consuming, for example, acquiring a complete mouse brain volume would require an estimated 3.6 million years per microscope under conventional settings. Preserving in situ 3D organization of cells within such samples of paramount importance for studies of tissue biology and pathological conditions. Leveraging a Hierarchical Attention Transformer (HAT) architecture, EMCellFiner achieves state-of-the-art super-resolution performance that substantially surpasses existing SOTA models. It effectively reconstructs intricate cellular ultrastructures across multiple scales and from diverse data sources obtained from different electron microscopy acquisition techniques, enabling 16-fold super-resolution with high fidelity. Crucially, its ability to recover structural details from heavily degraded inputs has profound practical implications. This capability drastically reduces imaging time and cost while maintaining or even enhancing image quality. This breakthrough opens avenues for more efficient experimental designs, particularly for large volume vEM studies.

In parallel, EMCellFound, built upon the MAE architecture, extends the utility of foundation models to high-level vision tasks. We demonstrate that EMCellFound can capture biologically meaningful features with a proficiency comparable to that of experienced cell biologists. It achieves state-of-the-art transferability in organelle recognition, detection, semantic, and instance segmentation tasks. Notably, it often matches or surpasses specialized models using only a fraction of the labeled data. Furthermore, the segmentation capabilities extend to 3D tasks, enabling efficient and accurate organelle reconstruction from 2D predictions. This approach exhibits superior performance compared to direct 3D segmentation, especially for irregularly structured organelles like the endoplasmic reticulum.

Although we have demonstrated that foundation models trained on vast EM datasets, such as EMCellFiner and EMCellFound, deliver substantial improvements in downstream task performance, their computational demands—particularly the reliance on large-scale datasets and high resource requirements—remain a practical limitation. Future optimizations focusing on few-shot learning, lightweight architecture, or employing token-based reconstruction objectives may mitigate these constraints. Furthermore, integrating large language models (LLMs) and generative models to establish structured memory for continual learning, and incorporating scientific knowledge, biological principles, and theories through machine learning tools for feature structuring offer promising avenues for further enhancing model adaptability and interpretability.

Finally, our models and dataset are being made publicly available to foster broader adoption and further innovation within the scientific community. We firmly believe that open science and data sharing are pivotal in accelerating interdisciplinary research that bridges biology, computer science, microscopy, and statistics. In summary, this work establishes a scalable and versatile foundation for the high-throughput, automated interpretation of vEM data. By significantly reducing the reliance on costly annotations and imaging time while delivering robust performance across species and tasks, the EMCFsys paves the way for large-scale exploration of life’s nanoscale architecture and accelerates discovery in cell biology.

## Methods

### Preparation of the Electron Microscopy Cell Foundation Dataset

To establish a comprehensive electron microscopy dataset for large-scale cellular representation learning, we systematically collected and curated a vast corpus of EM data (Fig. 1). The dataset integrates multi-source electron microscopy images from diverse origins, including public repositories (EMPIAR, BossDB, OpenOrganelle, and Nanotomy), datasets accompanying peer-reviewed publications, and shared archives from collaborating laboratories across China, the United States, Japan, and the Netherlands.

The collected data encompasses a broad spectrum of biological taxa, spanning animal, plant, and microbial specimens, and include both 2D and 3D imaging modalities. Multiple electron microscopy platforms were employed, such as TEM, SEM, FIB-TEM, FIB-SEM and so on. This diversity ensures comprehensive coverage of subcellular structures, including membranes, organelles, and fine ultrastructural details across a wide range of biological contexts.

### EMCFD Data Cleaning, Augmentation, and Quality Control

To construct a high-quality electron microscopy foundation dataset, we designed a multi-stage data processing pipeline encompassing slicing, cropping, augmentation, redundancy removal, and automated quality control (Supplementary Fig. 1). Distinct strategies were applied to 2D and 3D EM images to ensure both data diversity and structural integrity.

### 2D Image Processing

For 2D EM images, cropping was performed according to their native resolutions. Small-scale images from 512×512 to 4096×4096 pixels were partitioned into 1, 4, or 9 partially overlapping tiles, with a 20% overlap introduced to preserve contextual continuity across adjacent patches. Images exceeding 4096×4096 pixels were systematically cropped into 512, 768, 1024, 1536 pixels patches using a top-left to bottom-right scanning scheme. This process ensured balanced coverage of fine subcellular regions while avoiding boundary artifacts.

### 3D Image Processing

Given the high inter-slice redundancy inherent in vEM stacks, we adopted a sparse and multi-resolution slicing strategy. From each 3D stack, a subset of 2D slices (denoted as m frames) was uniformly sampled at fixed intervals to avoid oversampling highly correlated slices. Subsequently, we performed directional slicing along the X-Y, Y-Z, and X-Z planes to capture anisotropic structural information. Each extracted 2D slice larger than 5000×5000 pixels was cropped into image blocks of multiple scales, 512×512, 1024×1024, 1536×1536, 2048×2048, 3072×3072, and 4096×4096, following a left-to-right, top-to-bottom traversal pattern. To ensure non-redundancy, each original slice was processed only once per target size, and distinct slices were used for different resolutions. This strategy allowed the dataset to retain rich multiscale information while maintaining low inter-patch redundancy.

### Automated Quality Assessment

To further enhance dataset reliability, we trained a ResNet50^38^ binary classifier to distinguish high-quality EM images from those exhibiting noise, defocus, or mechanical artifacts. The classifier was trained on a manually labeled subset of 50,000 samples annotated by expert microscopists. Low-quality or corrupted patches were automatically discarded.

### Redundancy Removal and Similarity Filtering

A large-scale redundancy filtering pipeline was implemented to eliminate highly repetitive or overlapping image patches. We adopted a perceptual hashing based approach for efficient similarity detection. Each image was converted into a compact hash representation capturing its global visual structure and tonal distribution.

Pairwise Hamming distances between hashes were then computed, and image pairs with distances below a threshold of 5 were considered visually redundant. For each redundant group, only one representative image was retained, effectively reducing data duplication while preserving dataset diversity.

### Final Dataset Statistics

After all filtering and quality control stages, a total of 4,002,802 EM images of varying resolutions from 512×512 to 4096×4096 pixels were retained. The resulting dataset, EMCF dataset, represents one of the largest and most structurally diverse EM image corpora to date, encompassing multiscale, multi-organism, and multi-modality features suitable for large-scale foundation model pretraining.

### Image Restoration Model: EMCellFiner Design and Training

To reconstruct high-quality EM images from degraded observations, we developed EMCellFiner (Electron Microscopy Foundation Finer model), a transformer-based super-resolution foundation model built upon the HAT framework. The model learns a mapping function *F*: *L*−> *H*, where L and H denote the low-quality and high-quality EM image domains, respectively. The objective is to predict a high-fidelity reconstruction *F*(*L*) ≈ *H*, effectively restoring fine subcellular textures and structural integrity.

To simulate realistic degradation processes in EM imaging, we designed a two-stage stochastic degradation pipeline to generate the low-quality images *L* from the pristine high-resolution images *H* (sampled from the EMCF dataset). This degradation process aims to mimic various noise, blur, and compression artifacts commonly introduced during EM acquisition.

Each high-resolution image was first subjected to random upsampling and downsampling to simulate changes in magnification and scanning area.

### Stage1

A random average blur with kernel size in the range from 1 to 21 was then applied to imitate optical defocus and beam-induced diffusion. Subsequently, the image was downsampled using one of three randomly selected interpolation modes, including bicubic, bilinear, and area sampling, followed by the addition of Gaussian noise range 1 to 30, Poisson noise, and JPEG compression artifacts.

### Stage2

A second round of random blurring with the kernel from 1 to 21 and then 4× downsampling was performed to achieve the target degradation level, followed by an additional Gaussian noise injection. This two-step degradation ensures that the generated images encapsulate a wide spectrum of realistic low-quality conditions observed in diverse EM imaging setups.

### Model Architecture and Training Strategy

Following the configuration of HAT, EMCellFiner adopts a hierarchical transformer architecture with hybrid attention mechanisms for spatial and channel-wise feature refinement.

We employed a GAN-based training scheme, where EMCellFiner acts as the generator, and a lightweight CNN-based discriminator (VGG-19) guides perceptual realism.

The overall objective function combines pixel-level reconstruction loss, perceptual loss, and adversarial loss:

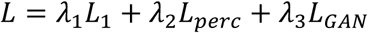

where *L*_1_ denotes the L1 loss between the restored and ground-truth images, *L_perc_* is the perceptual loss computed on VGG feature space, and *L_GAN_* corresponds to a vanilla adversarial loss encouraging photorealistic reconstruction.

Training was conducted on 8 NVIDIA H800 GPUs for 400,000 iterations with a batch size of 16. The entire process took approximately 4 days. Adam optimizer was used with an initial learning rate of 2×10^-4^, decayed following a cosine annealing schedule. All images were randomly cropped into 64×64 patches as input, and the model was trained to reconstruct corresponding 256×256 high-resolution outputs.

### EMCellFound architecture and implementation

EMCellFound was an encoder pretrained within a masked autoencoding framework consisting of a paired encoder-decoder architecture. To achieve both scalability and computational efficiency, a 12-layer Vision Transformer (ViT) encoder was adopted to model long-range dependencies and capture fine structural representations typical of EM imagery. A lightweight 9-layer Transformer served as the decoder, with an embedding dimension of 512, responsible for reconstructing the masked portions of the input image. This asymmetric encoder–decoder design balances representation capacity and training efficiency, allowing EMCellFound to learn high-level semantic priors while maintaining feasible training costs for large-scale EM datasets.

Following the training paradigm of MAE, 75 % of the image patches were randomly masked during pretraining. The model was then optimized to reconstruct the original image from the remaining 25 % of visible patches, enabling it to learn high-level semantic structures from partial visual cues. We adopted the default masking ratio suggested in the MAE paper. All model parameters were initialized randomly prior to training.

Optimization was performed using the AdamW optimizer with an initial learning rate of 0.0002, scheduled via a warmup-cosine decay strategy. The learning rate was linearly warmed up from 1×10^-7^ to the base rate during the first 20 epochs. Training was conducted for 1500 epochs with a batch size of 512, using mixed-precision (FP16) to accelerate convergence and reduce memory usage. The final checkpoint from the last epoch was retained as the pretrained foundation model for downstream finetuning and adaptation. Training was performed on 8 NVIDIA H800 GPUs and completed in approximately 3 days.

Standard loss and optimization objectives were employed following MAE conventions. The training loss is defined as a mean squared error between the reconstructed pixels and the original image, computed only over masked patches:

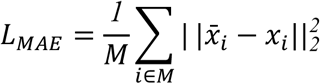

where M denotes the set of masked patches, xi is the original pixel value of patch i, and x-i represents the reconstructed pixel value predicted by the decoder. By restricting the reconstruction loss to the masked regions, the encoder is encouraged to learn semantic and structural priors rather than trivial low-level pixel statistics, a property particularly advantageous for modeling the fine-grained morphology and texture diversity inherent in EM images.

For comparative evaluation, we trained additional self-supervised baselines, DINO, MoCov3, and All4One, under identical ViT backbone configurations and hyperparameter settings. This comparison allowed us to quantitatively assess the effectiveness of the MAE-based pretraining strategy for EM imagery representation learning.

### Adaptation to zero-shot classify

To evaluate the generalization capability of the EMCellFound model on downstream tasks, we designed a zero-shot classification pipeline that requires no task-specific fine-tuning. In this setup, only the encoder component of EMCellFound was retained, and extract the feature embedding from enconder output. For classification, we adopted a non-parametric K-Nearest Neighbors classifier to directly perform category prediction based on the extracted embedding. Each test image x was passed through the frozen encoder to obtain its latent representation f(x) ∈ ℝ^d^.

The predicted label y was then determined according to the majority label among its k nearest neighbors in the feature space:

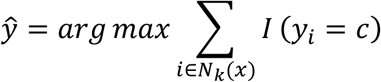

where N_k_(x) denotes the set of k nearest neighbors of image x, y_i_ is the class label of neighbor i, and C is the set of all categories.

The neighborhood was defined according to cosine distance, ensuring that similarity was determined by feature orientation rather than magnitude differences. The number of neighbors *k* was set as 5, which empirically balanced local consistency and global separability across categories.

No gradient update or parameter optimization was performed on the encoder during this process, ensuring a pure zero-shot evaluation of its representation quality. The classification accuracy was computed across all organelle categories to assess the encoder’s intrinsic discriminative ability and semantic separability in feature space, without any supervision or task-specific adaptation

### Adaption to semantic segmentation

To enable effective adaptation of EMCellFound to downstream semantic segmentation tasks, we integrated a segmentation head on top of the pretrained encoder to achieve pixel-level segmentation. Given its strong performance in prior segmentation studies, we adopted UPerNet as the segmentation head to enhance both accuracy and generalization capability. The overall architecture and parameter configuration of UPerNet are illustrated in Supplementary Fig. 19. Its final layer acts as a pixel-wise classifier that outputs the predicted organelle category for each pixel. Between the pretrained ViT encoder and the UPerNet segmentation head, we inserted a lightweight neck module designed to aggregate and fuse multi-scale feature representations from both shallow and deep layers. This fusion facilitates richer contextual understanding of fine structural variations in EM images. The overall objective function combines Dice Loss and Cross-Entropy Loss, formulated as

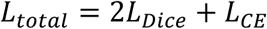

where *L_Dice_* is the dice loss and *L_CE_* is the cross entropy loss.

All segmentation experiments were conducted under unified training configurations to ensure performance comparability across models.

### Adaption to instance segmentation

To enable EMCellFound to support instance segmentation tasks, we appended a lightweight RTMHead to the pretrained encoder. The RTMHead module provides an efficient balance between inference speed and segmentation accuracy, making it well suited for large-scale EM datasets with densely distributed organelles. Following the standard configuration, the RTMHead was trained jointly with fine-tuning of the EMCellFound encoder to adapt its representations toward instance-level feature discrimination. The training objective consisted of three components: a classification loss, a bounding box regression loss, and a mask segmentation loss, formulated respectively as:

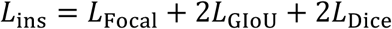

where *L*_Focal_ denotes the Quality Focal Loss, which improves classification confidence calibration and positive-negative sample balance; *L*_GIoU_ is the Generalized IoU (GIoU) Loss, measuring spatial overlap quality between predicted and ground-truth bounding boxes; *L*_Dice_ corresponds to the Dice Loss, emphasizing accurate mask prediction for small and irregular organelle instances.

This composite loss design ensures that EMCellFound not only preserves global contextual understanding from pretraining but also acquires fine-grained instance-level discrimination required for EM image analysis.

### Adaption to 3D image stack segmentation

To extend the capability of EMCellFound to 3D cellular structure analysis, an efficient 2D slice based 3D segmentation and reconstruction pipeline was developed. This pipeline leverages the strong representational capacity of the pretrained encoder to achieve high-accuracy volumetric segmentation under extremely low annotation cost. The 3D reconstruction process begins with precise cross-slice segmentation. Specifically, semantic or instance segmentation is performed on 3D image stacks by decomposing them into a sequence of consecutive 2D slices. The pretrained encoder and the corresponding segmentation heads, identical to those used in 2D tasks, are independently applied to each slice. To evaluate the few-shot adaptability of EMCellFound, the pretrained encoder and segmentation head are fine-tuned using only a very small subset of manually annotated slices. From a 3D stack containing over N (>1000) consecutive sections, only 1-10 slices (approximately 0.1%-1% of the stack) are randomly selected and annotated. The model is then fine-tuned on these sparse annotations to generalize toward accurate segmentation on unannotated slices.

After segmentation, the 3D volume is reconstructed by re-stacking all predicted 2D segmentation masks back into their original spatial positions, yielding a voxel-wise 3D prediction. To address potential inter-slice inconsistencies introduced by independent 2D predictions, lightweight voxel-level post-processing, including 3D morphological operations and connected component analysis, is applied to ensure structural smoothness and cross-slice continuity. The final reconstructed 3D volumes are subsequently used for quantitative organelle analysis (e.g., volume and surface area estimation) and visualization, such as volumetric rendering or surface mesh reconstruction.

## Data and code availability

The EMCF dataset for the current study is available in the Science Data Bank repository at https://cstr.cn/31253.11.sciencedb.32271. The EMCFsys plugin of napari is available on Github, at https://github.com/yzy0102/emcfsys.

## Acknowledgements

This work was supported by the National Natural Science Foundation of China (32572179). We sincerely thank Hangzhou Chengfengerlai Digital Technology Co., Ltd’s support. We also thank Ben Giepmans of Universitair Medisch Centrum Groningen for the data assistance and feedback.

## Contributions

\# Zeyu Yu, Jiansheng Guo, Feng Liu contributed equally to this work.

Zeyu Yu, Xuping Feng and Mengze Du conceived the study and designed the experiments. Zeyu Yu, Jiansheng Guo, Feng Liu, Mengze Du, Shan Xu, Guowei Zhang and Li Xie performed data acquisition and curation. Gaoliang Deng, Chen Rui. provided essential computational resources and storage infrastructure. Zeyu Yu, and Xuping Feng wrote the manuscript. Yong He and Bo Han reviewed and edited the manuscript.

