## Supplement Information for "EMCF ecosystem: Towards pretrained foundation model for electron microscopy image analysis"

### Supplementary Information

***This file contains Supplementary Methods, Supplementary Figures, Supplementary Tables and Supplementary References.***

#### Supplementary Figures

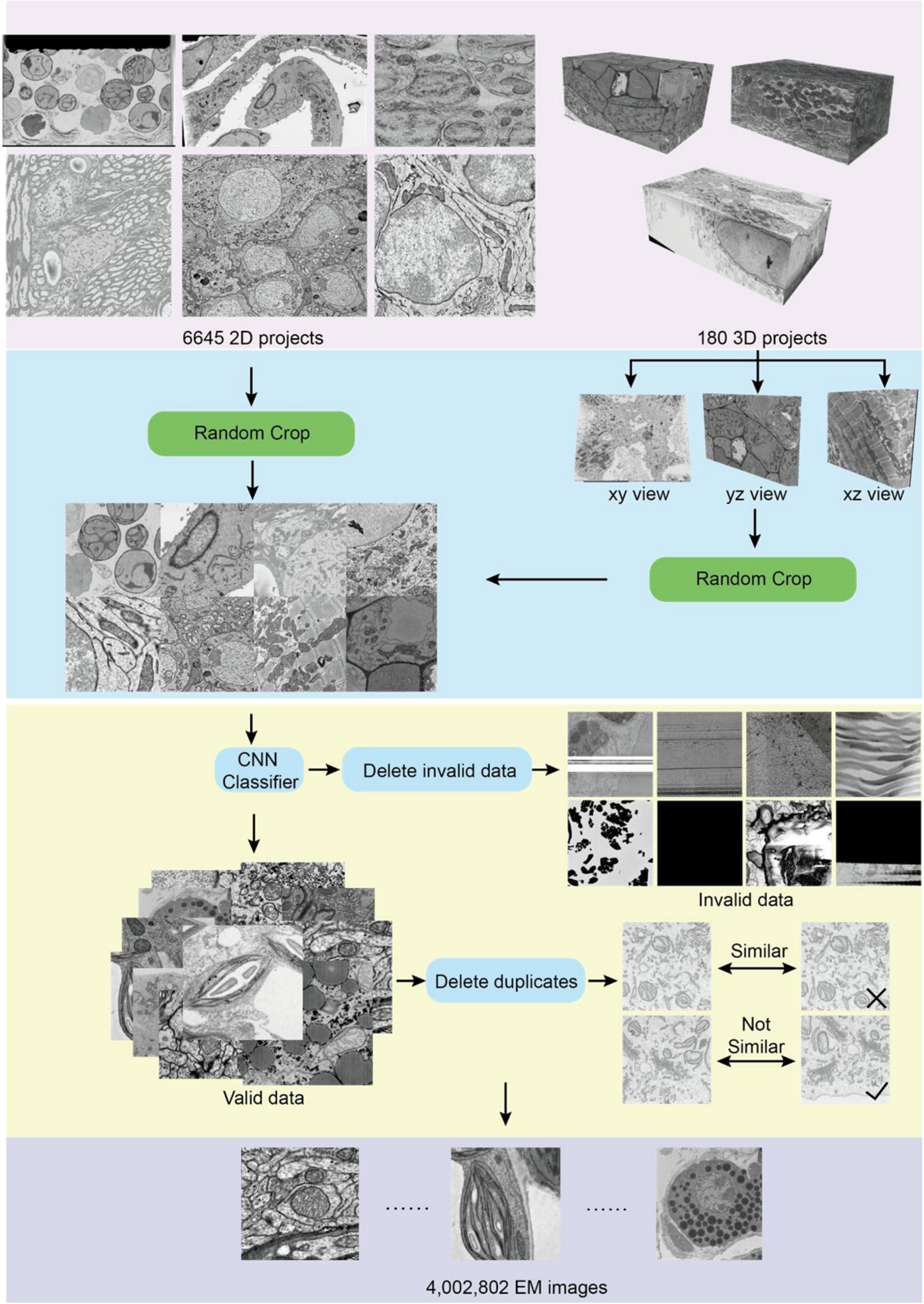

**Supplementary Fig. 1.** Image processing workflow for preparing the high-quality Electron Microscopy Cell Foundation (EMCF) dataset.

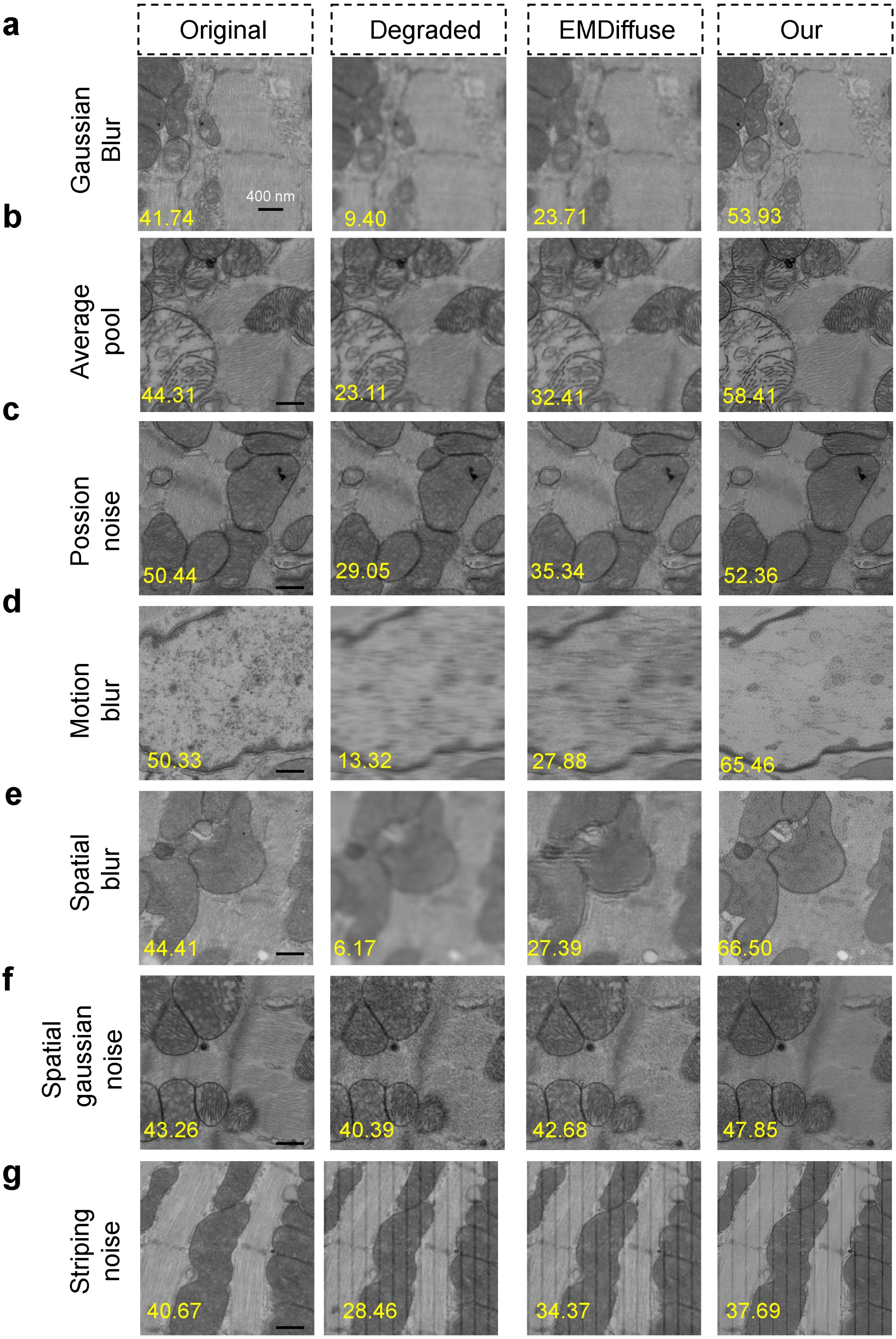

**Supplementary Fig. 2** Comparison of SR results from EMDiffuse and EMCellFiner on microscopy images degraded via different synthetic perturbations. a, 3×3 Gaussian blur. b, 4×4 mean filtering. c, Poisson noise (intensity-dependent, simulating low-light conditions). d, Motion blur (9-pixel horizontal kernel). e, 5×5 mean filtering (detail suppression). f, Additive white noise (σ=10, per-pixel i.i.d.). g, Column-wise stripe artifacts (10-pixel spacing, 20-intensity reduction, mimicking EM scan noise). Degraded inputs were generated by 4× downsampling with combined perturbations to emulate real-world EM image distortions. Sharpness scores are shown for related images.

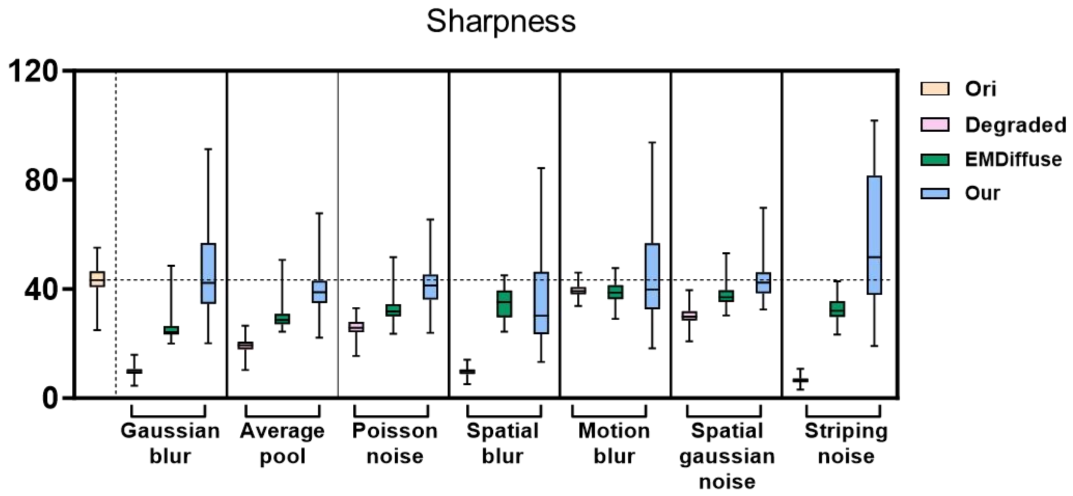

**Supplementary Fig. 3** Comparison of sharpness for SR reconstructions generated by EMCellFiner and EMDiffuse across various degradation patterns. The data sources used for these tests are detailed in Supplementary Table 1.

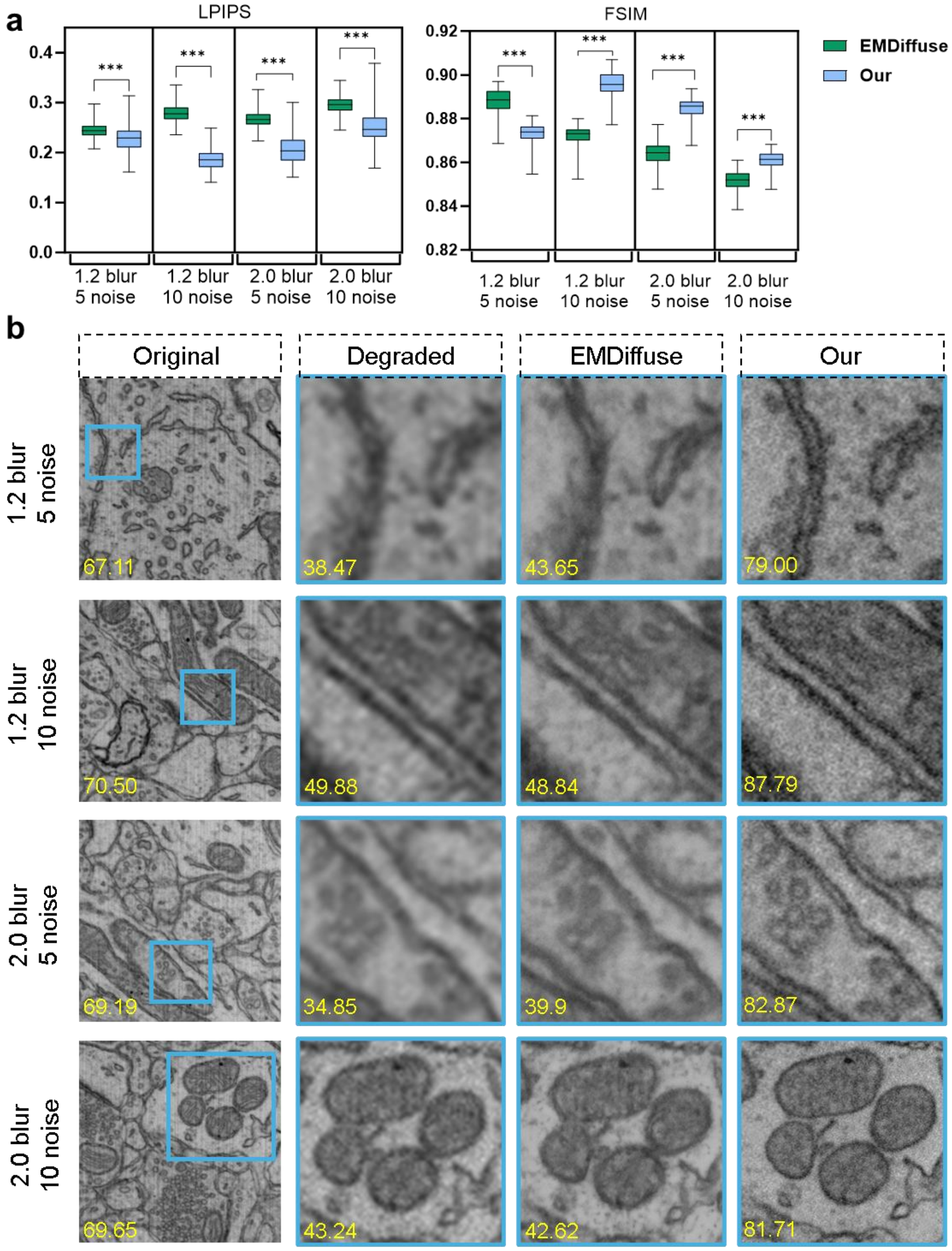

**Supplementary Fig. 4** Comparison super-resolution reconstructions generated by EMCellFiner and EMDiffuse under different blur and noise. a, Statistic comparison on the dataset (Supplementary table 1) in terms of LPIPS and FSIM. b, Representative SR images reconstructed by EMCellFiner and EMDiffuse. Sharpness values are shown for related images.

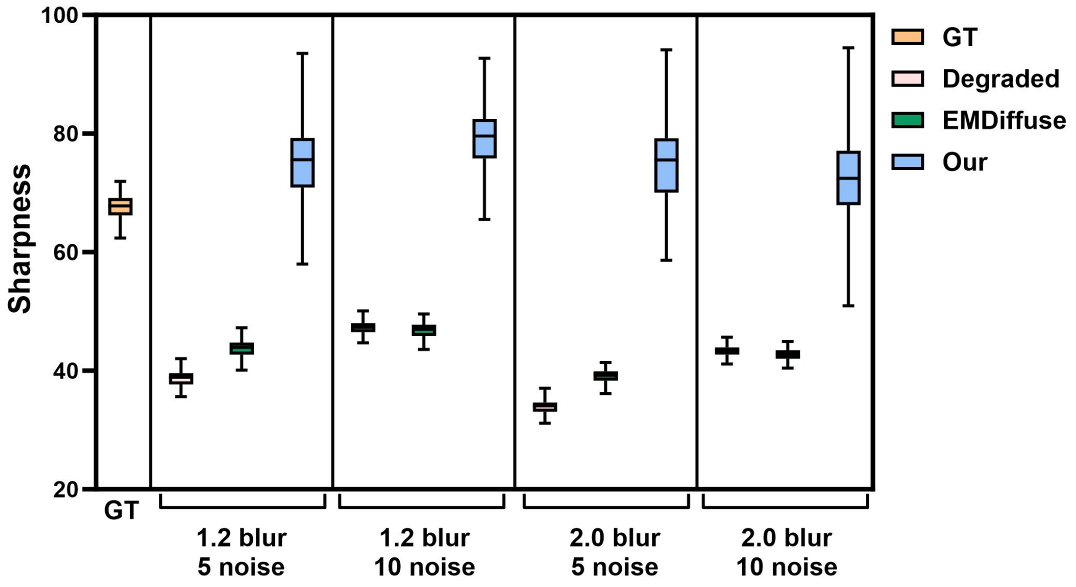

**Supplementary Fig. 5** Sharpness comparison super-resolution reconstructions generated by EMCellFiner and EMDiffuse under different blur and noise. The data sources used for these tests are detailed in Supplementary Table 1.

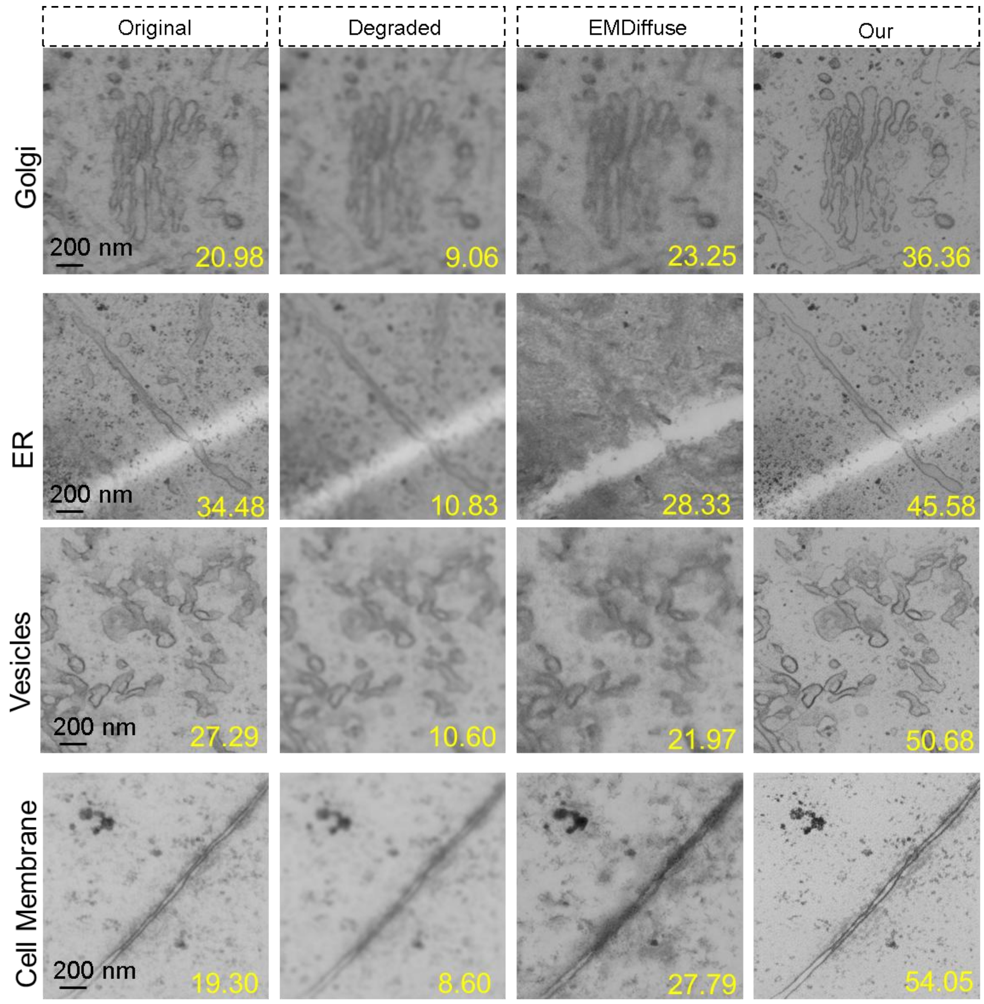

**Supplementary Fig. 6** Comparative analysis of super-resolution performance on the different organelles from Kuiper2022NCB dataset by EMDiffuse and EMCellFiner. Degraded images were generated from ground truth images (512px) via 3×3 Gaussian blurring and 4× downsampling.Sharpness values are shown for related images.

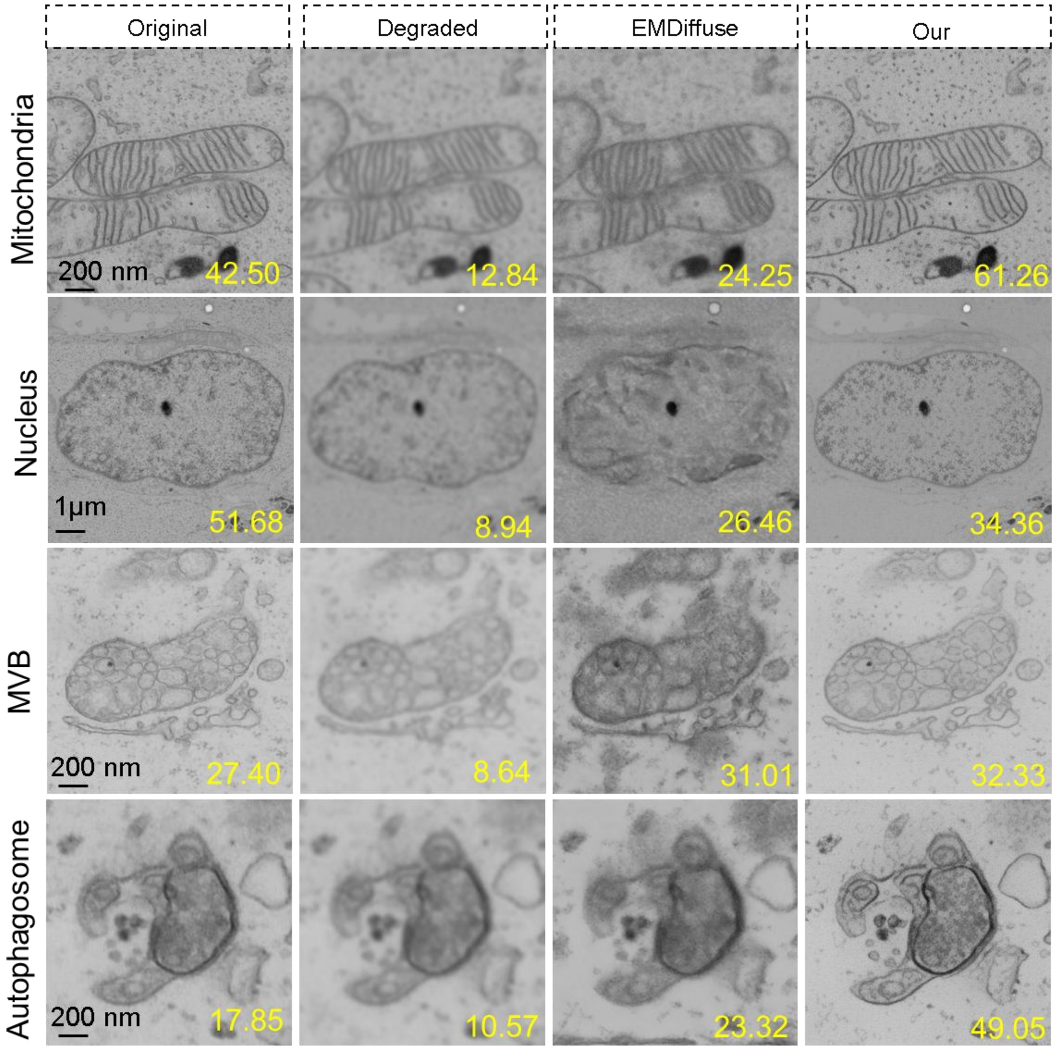

**Supplementary Fig. 7** Comparative analysis of super-resolution performance on the different organelles from Kuiper2022NCB dataset (ref) by EMDiffuse and EMCellFiner. Degraded images were generated from ground truth images (512px) via 3×3 Gaussian blurring and 4× downsampling.Sharpness values are shown for related images.

As in Supplementary Fig. 7, for each organelle type shown, EMCellFiner reconstructions consistently exhibit superior structural clarity compared to those from EMDiffuse. This visual improvement is quantitatively supported by the sharpness values displayed above each image. Notably, the sharpness scores for EMCellFiner reconstructions frequently exceed those of the original ground truth images, suggesting that our method not only restores but actively enhances high-frequency features, leading to a perceptually sharper output.

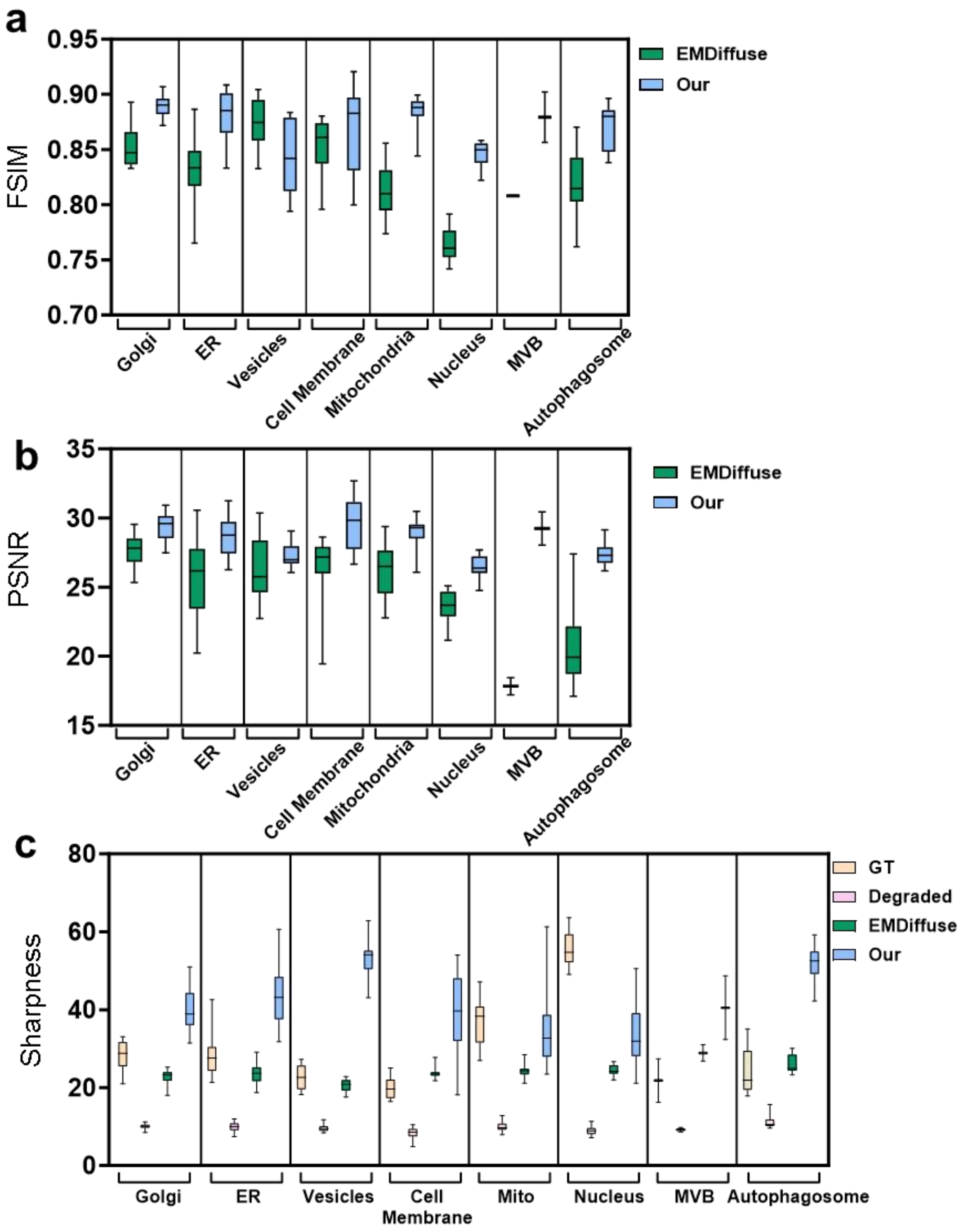

**Supplementary Fig. 8** Comparison of FSIM , PSNR and sharpness for super-resolution reconstructions generated by EMCellFiner and EMDiffuse under high Signal-to-Noise Ratio conditions. Degraded images were generated from ground truth images (512px) via 3×3 Gaussian blurring and 4× downsampling.The data sources used for these tests are detailed in Supplementary Table 1.

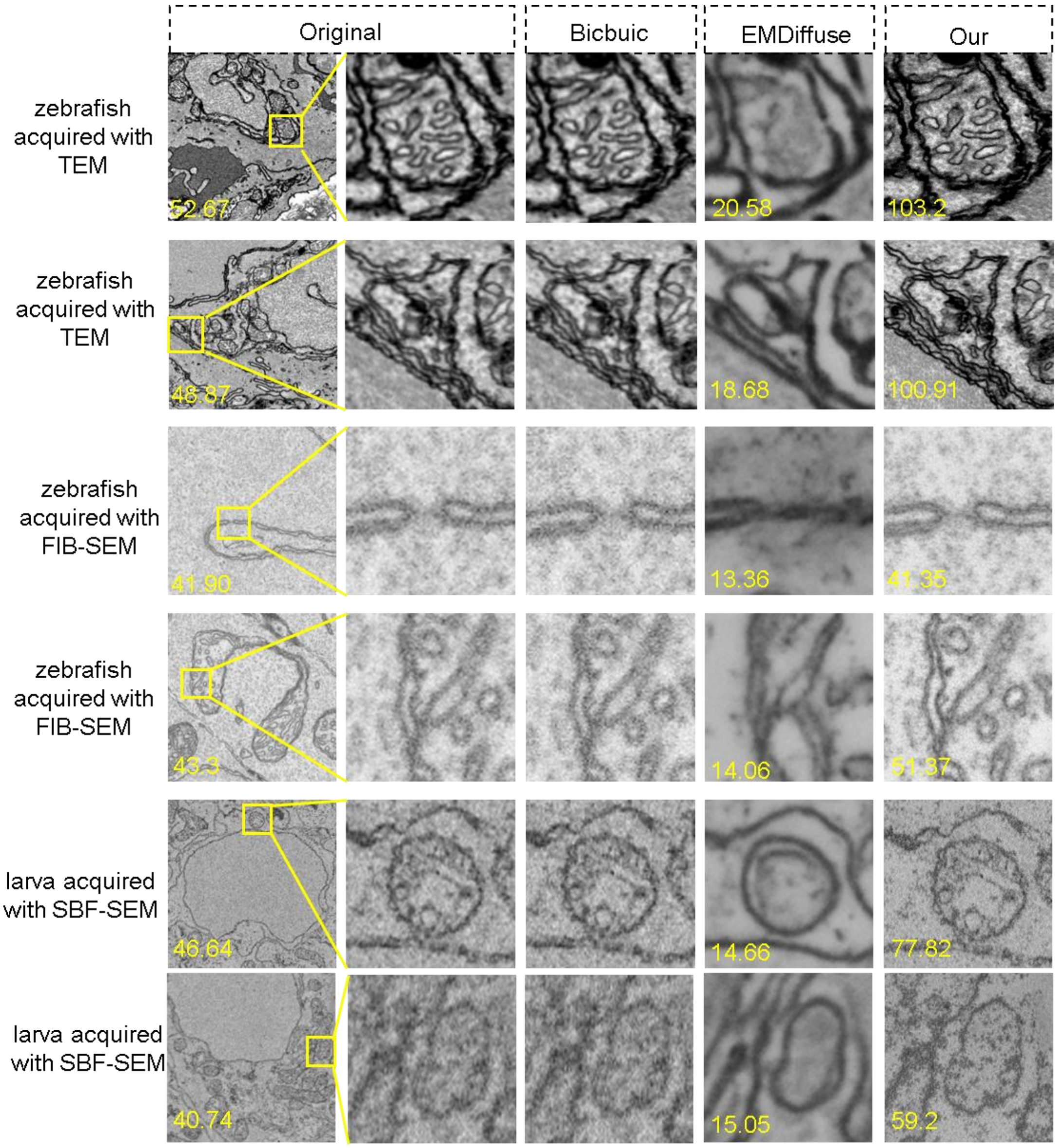

**Supplementary Fig. 9** Representative SR images reconstructed by EMCellFiner and EMDiffuse under low Signal-to-Noise Ratio conditions. Sharpness values are shown for related images. The data sources obtained from different electron microscopy acquisition used for these tests are detailed in Supplementary Table 1.

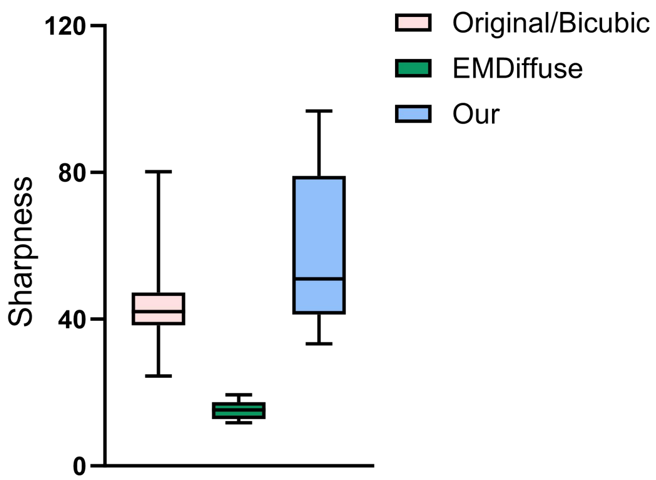

**Supplementary Fig. 10** Sharpness comparison SR reconstructions generated by EMCellFiner and EMDiffuse under low Signal-to-Noise Ratio conditions. The data sources obtained from different electron microscopy acquisition used for these tests are detailed in Supplementary Table 1.

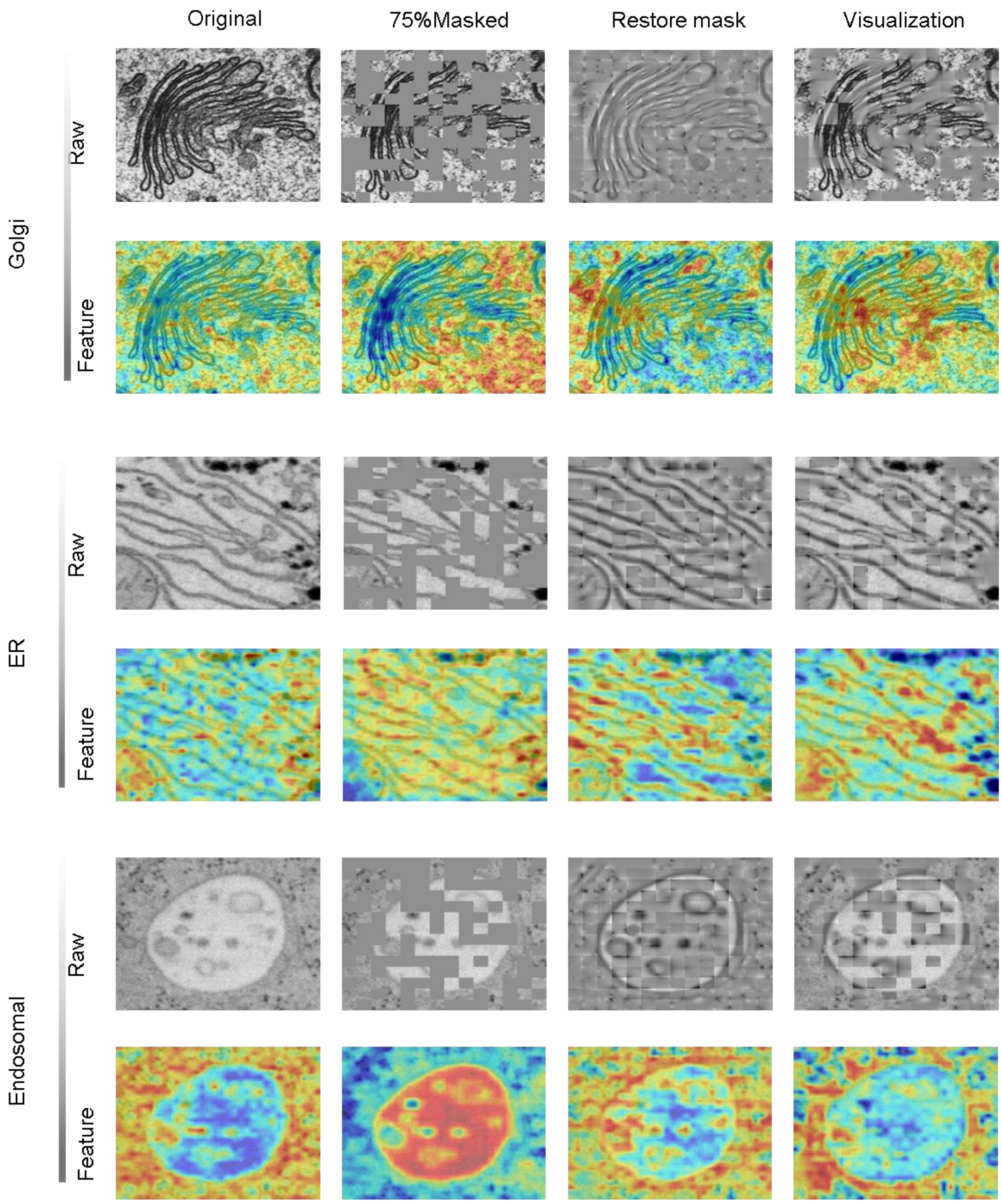

**Supplementary Fig. 11** Visualization of CAMs from different layers of EMCellFound for identifying Golgi apparatus, endoplasmic reticulum (ER), and endosomes. Upper row: Input image processing pipeline (raw image → 75% masked input → reconstructed feature map → final reconstruction). Bottom row: Attention heatmaps from Transformer layers 3, 6, 9, and 12, highlighting maximal attention regions.

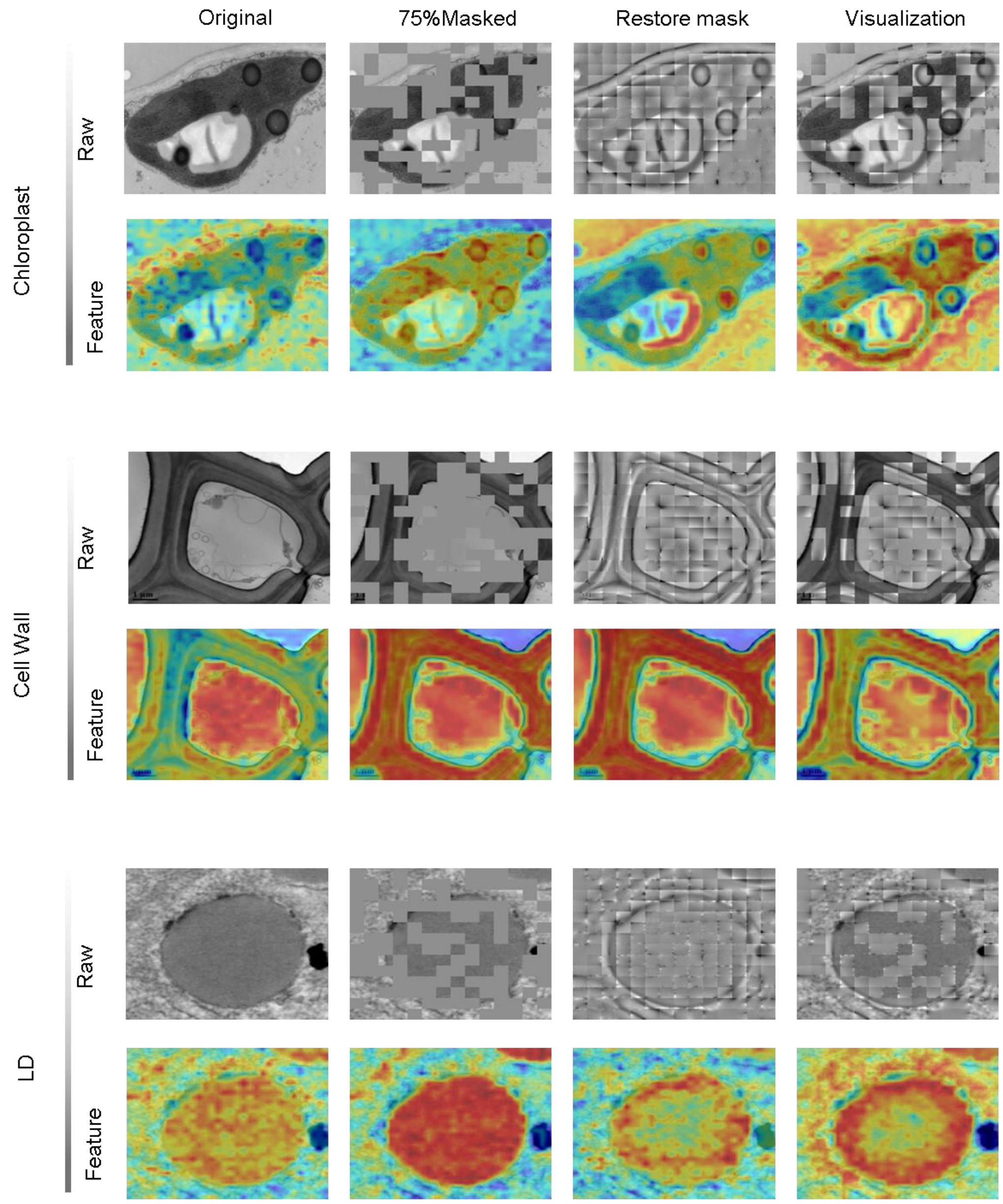

**Supplementary Fig. 12** Visualization of CAMs from different layers of EMCellFound for identifying chloroplast, cell wall, Lipid droplets (LD) .Upper row: Input image processing pipeline (raw image → 75% masked input → reconstructed feature map → final reconstruction). Bottom row: Attention heatmaps from Transformer layers 3, 6, 9, and 12, highlighting maximal attention regions.

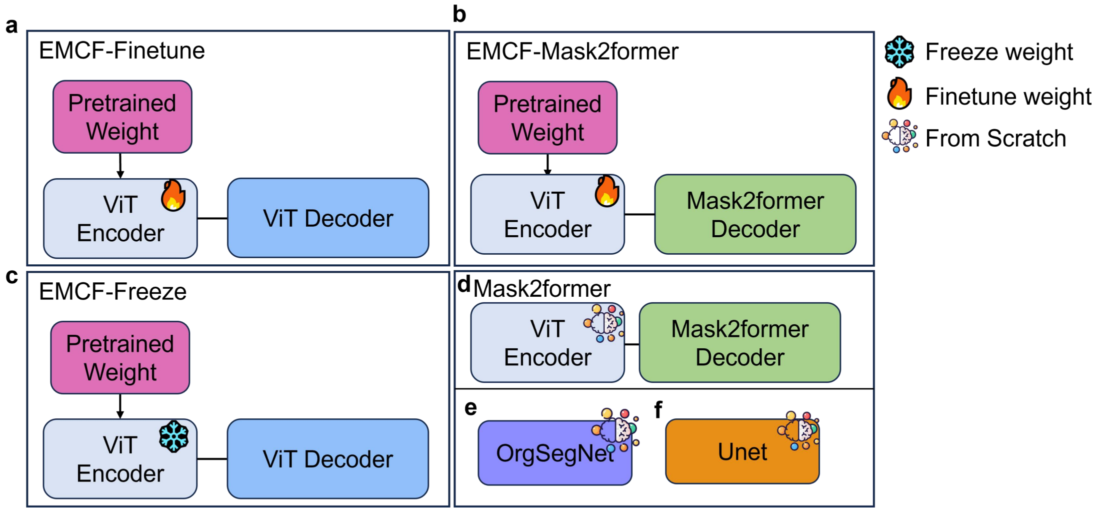

**Supplementary Fig. 13** Model implementation and training strategies for comparative evaluation in Fig. 4. a, EMCF-Finetune. The pretrained EMCellFound (EMCF) encoder (a Vision Transformer (ViT) architecture) is integrated with a ViT-based decoder as the segmentation head. All model parameters (encoder and decoder) are fine-tuned end-to-end for semantic segmentation. b, EMCF+Mask2Former. This strategy employs a hybrid architecture where the pretrained EMCF (ViT Encoder) is coupled with a Mask2Former segmentation head. Both the pretrained encoder and the Mask2Former head parameters are fine-tuned. c, EMCF-Freeze. The pretrained EMCF (ViT Encoder) parameters are kept fixed (frozen), and only the parameters of the ViT-based decoder (segmentation head) are trained. d, Mask2Former (Baseline). A standard Mask2Former segmentation model is trained from scratch. Both its internal ViT Encoder and Mask2Former Decoder parameters are randomly initialized before training. e, OrgSegNet and UNet (Baselines). The specialized domain model OrgSegNet and the general baseline UNet model are trained from scratch. Their parameters are randomly initialized prior to training.

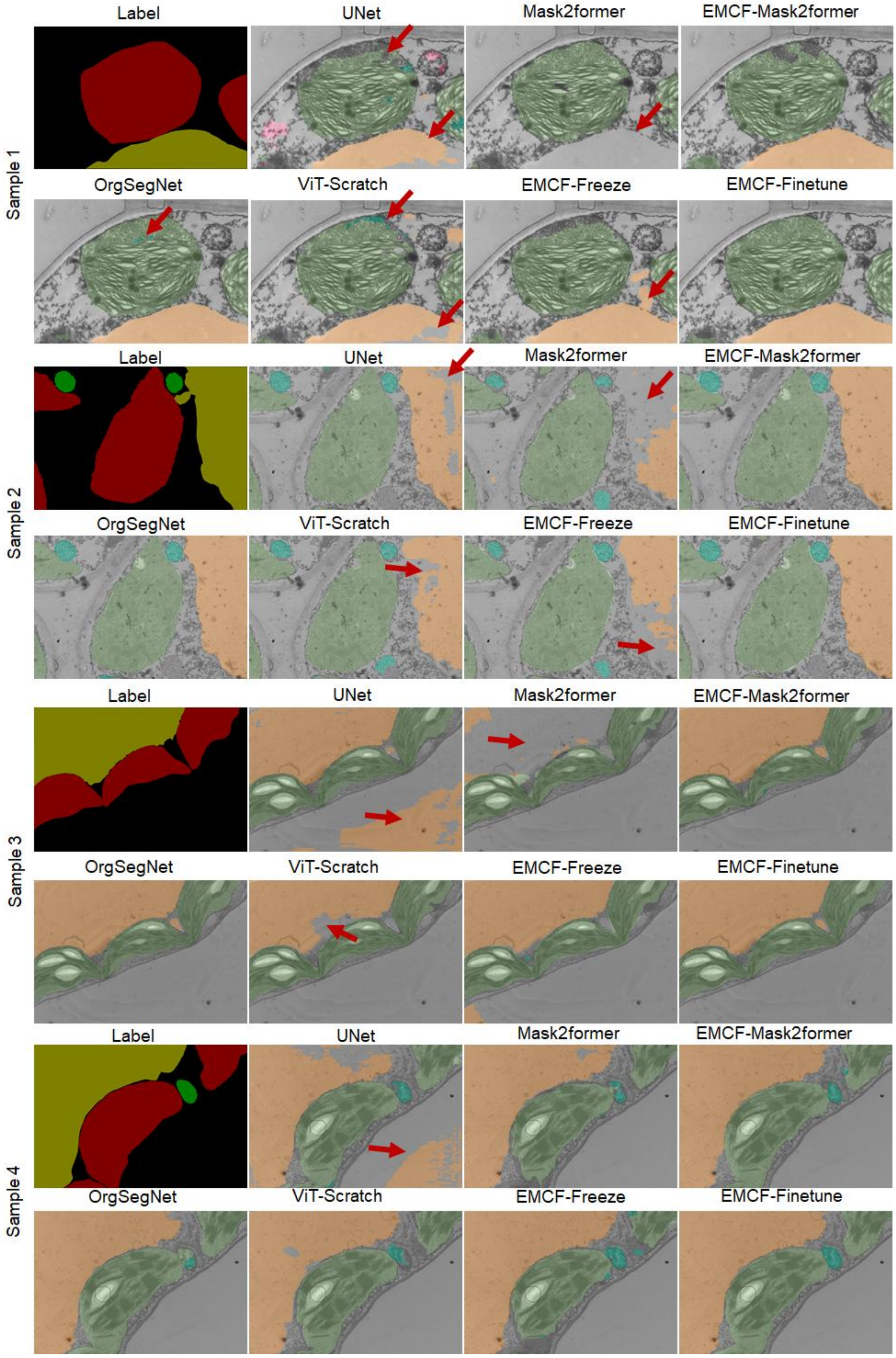

**Supplementary Fig. 14** Representative TEM images of the diverse morphology of organelles segmented by seven models. The red arrows indicate recognition errors.

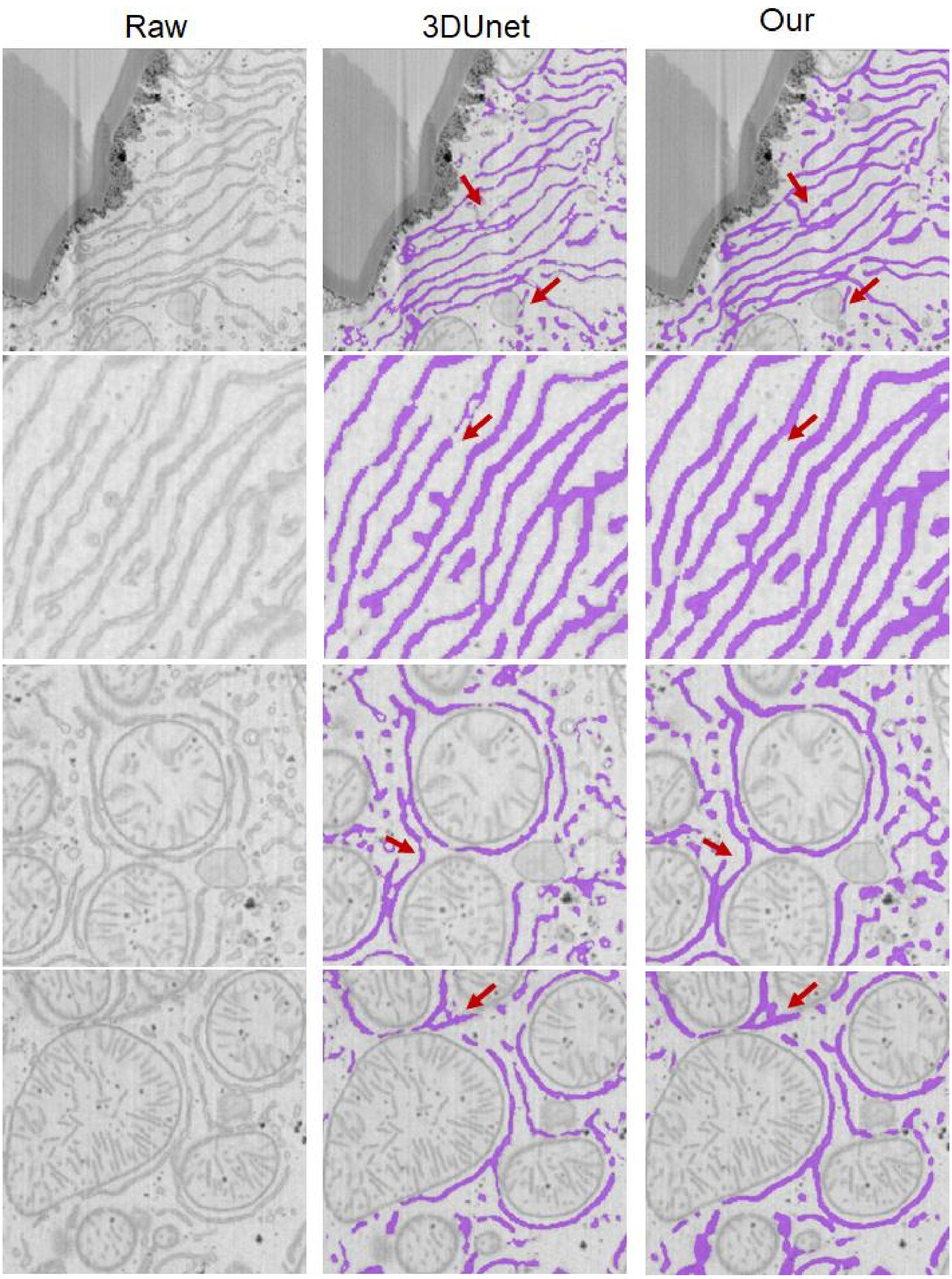

**Supplementary Fig. 15**. Comparison of ER segmentation results between 3D U-Net and EMCF on the Liver-6 dataset. EMCF achieves more continuous ER segmentation in 2D EM images, while 3D U-Net generates fragmented ER structures, leading to discontinuities in the 3D reconstruction. Red arrows highlight the differences between the two methods.

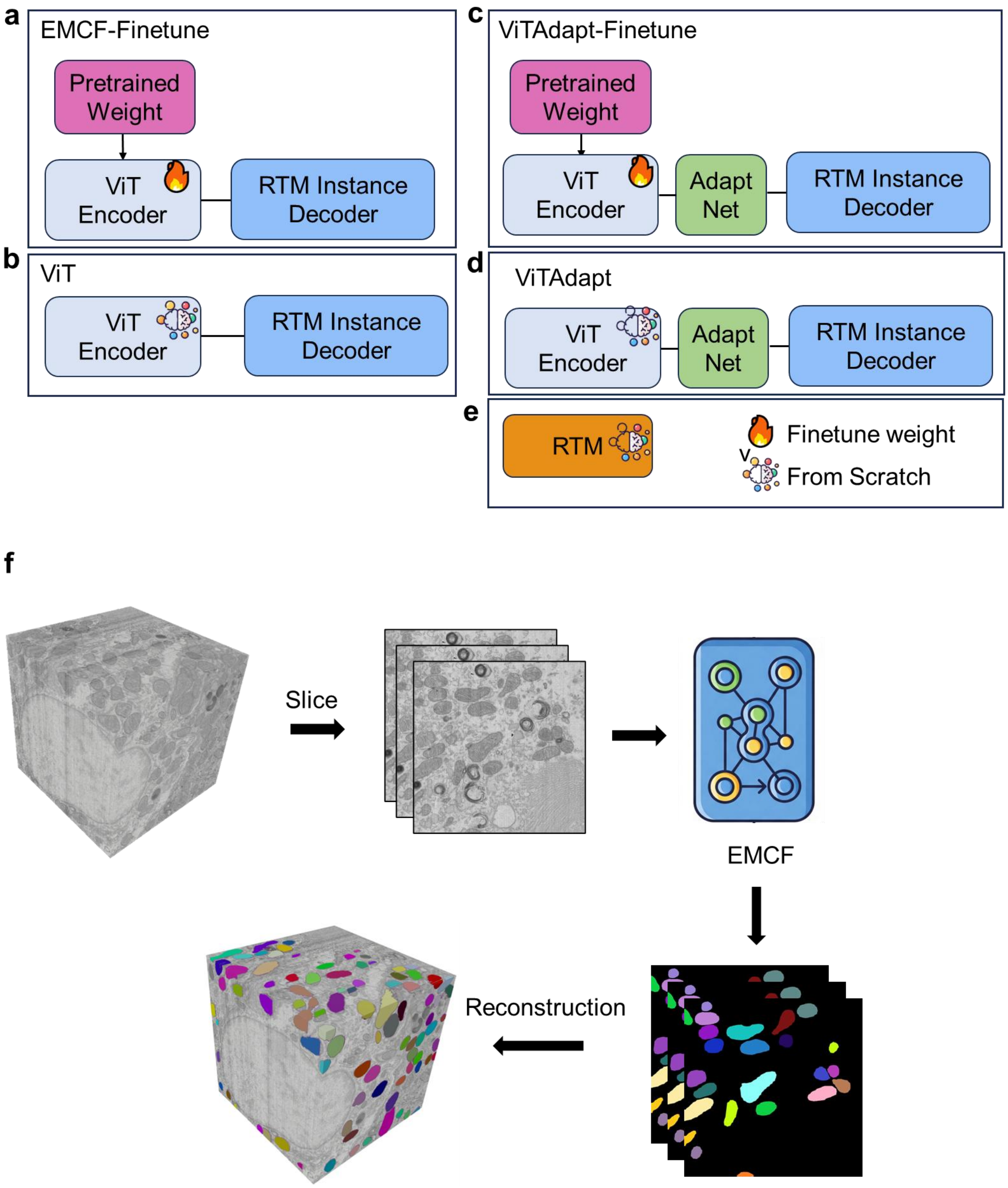

**Supplementary Fig. 16**. Model implementation and training strategies for comparative evaluation in Fig. 5. a, EMCF-Finetune. The pretrained EMCellFound (EMCF) encoder (a Vision Transformer (ViT) architecture) is integrated with a RTM decoder as the instance head. All model parameters (encoder and decoder) are fine-tuned end-to-end for instance segmentation. b, ViT . The ViT Encoder with RTM Instance Decoder parameters are initiate from scratch. c, ViTAdapt-Finetune. The pretrained EMCellFound (EMCF) encoder (a Vision Transformer (ViT) architecture) with the Adapt structure is integrated with a RTM decoder as the instance head. All model parameters (encoder and decoder) are fine-tuned end-to-end for instance segmentation. d, ViT-Adapt (Baseline). The ViT Encoder with Adapt structure and RTM Instance Decoder parameters are initiate from scratch. e. RTM. The RTM model with CNN backbone, whose parameters are initiate from scratch. f. the pipline of Mitochondria 3D reconstruction using EMCF model.

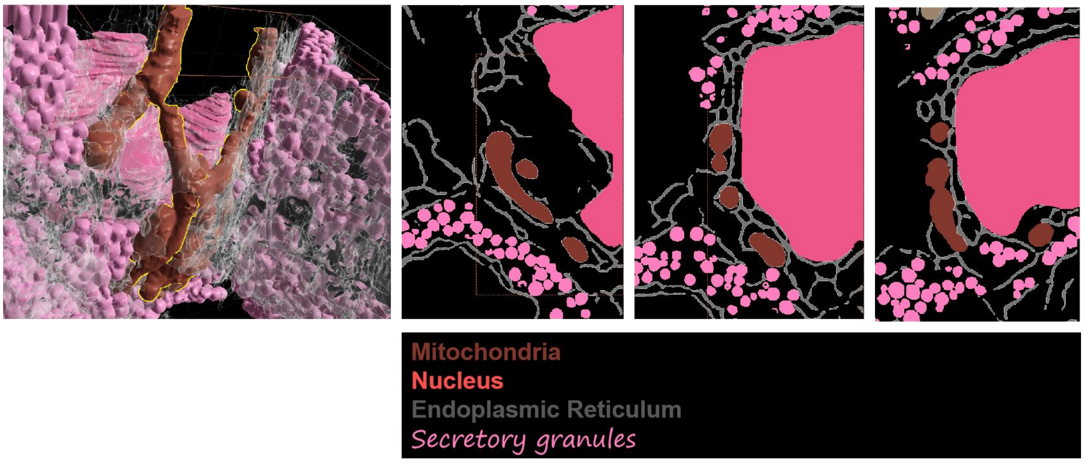

176 **Supplementary Fig. 17.** Representative zoomed-in 3D models of anterior pituitary
177 cells.

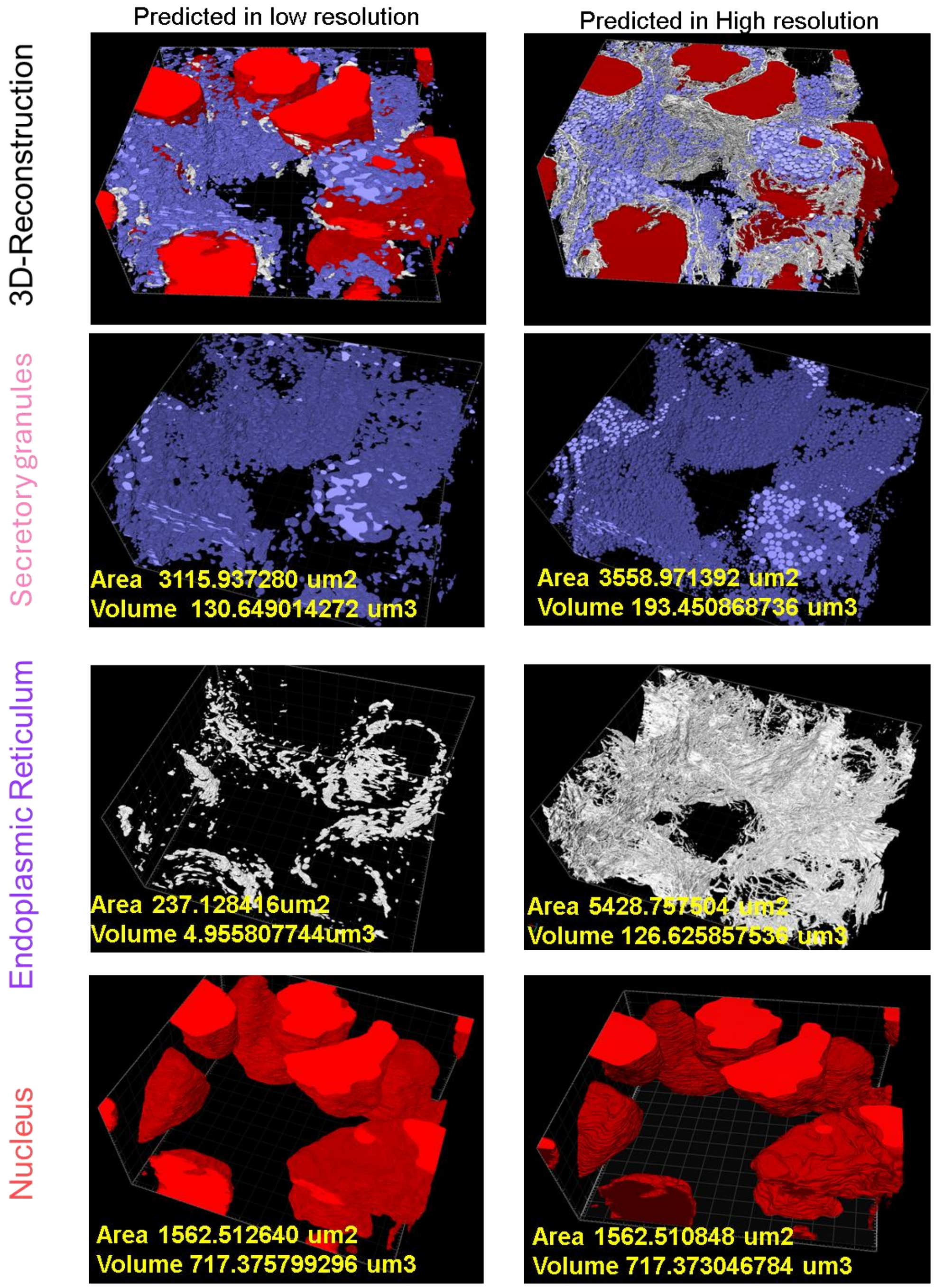

**Supplementary Fig. 18.** Comparative 3D reconstruction of four organelle types, showcasing the low-resolution input and the high-resolution output achieved by EMCellfiner.

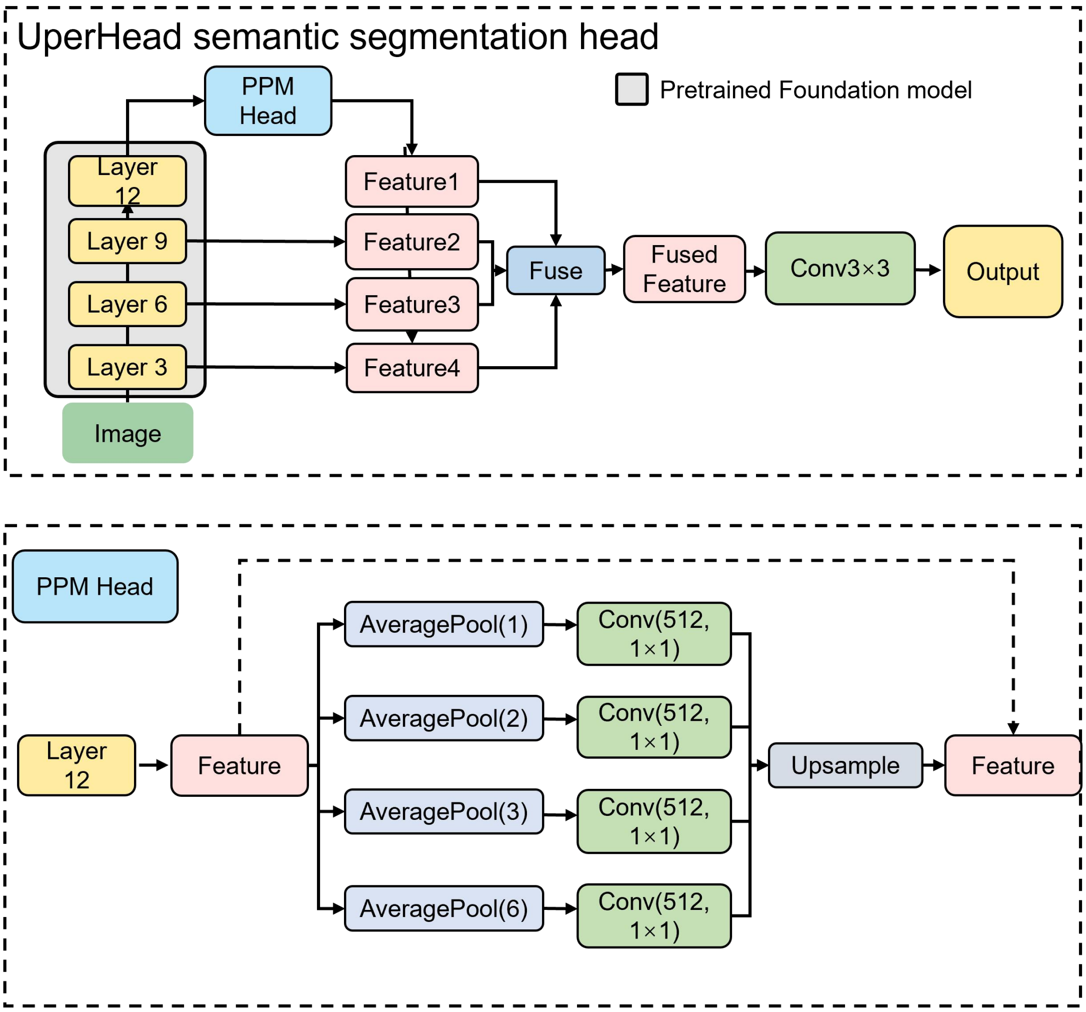

**Supplementary Fig. 19** UperNet Head for segmentation for EMCellFound backbone.

**Supplementary Table 1 Information on datasets involved in predicting.**

| **Fig** | **dataset** | **dataset size** | **data description** | **data link** |
|---|---|---|---|---|
| **fig2c\d supple fig3、4** | nijholt | 117 | / | https://www.nanotomy.org/ |
| **supple fig5、6** | jrc_mus-nacc-2 | 100 | Nucleus accumbens from a wild-type, adult male mouse | https://www.openorganelle.net/datasets/jrc_mus-nacc-2 |
| **supple fig7、8\9** | Kuiper2022NCB | golgi-14<br>er-18<br>vesicles-11<br>cell membrane-14<br>mito-14<br>nucleus-17<br>mvb-2<br>autophagosome-13 | / | https://www.nanotomy.org/ |
| **fig2e supple fig10\11** | EMPIAR-10459\Hiroki Nishida2021\Zebrafish_Lobe | 35 | / | https://empiar.ipr.pdbj.org/en/entry/10459/ |
| **supple fig 15** | Plant TEM dataset | 650 images for training<br>180 images for validation<br>180 images for testing<br>180 | / | plantorganelle hunter dataset in https://cstr.cn/31253.11.sciencedb.01335 |
| **supple fig 15** | jrc-liver-6 | 3<br>2000*2000 pixels images for training | Liver cells from male mouse 10 weeks old | https://www.openorganelle.net/datasets/jrc_mus-liver-6 |

| | | | | |
|---|---|---|---|---|
| | | volume size:8000*8000*8500 | | |
| **fig5ab** | LEE | training:133<br>validation:88<br>test:89<br>1024*1024 pixels images | Electron Microscopy data used in a study of an excitatory network in Mouse V1. | https://bossdb.org/project/lee2016 |
| **fig5ab** | jrc_mus-kidney-2 | train:42<br>test:30<br>val:38<br>1024*1024 pixels images | kidney cells from wild-type, 8 week old mouse | https://www.openorganelle.net/datasets/jrc_mus-kidney-2 |
| **fig5ab** | bock | train:87<br>test:30<br>val:30<br><br>1024*1024 pixels images | Volume of mouse primary visual cortical data | https://bossdb.org/project/bock2011 |
| **fig5cde** | jrc_mus-kidney-2 | 1024*1024*1024 randomly selected vloume data | kidney cells from wild-type, 8 week old mouse | https://www.openorganelle.net/datasets/jrc_mus-kidney-2 |
| **fig6-c** | EM image stacks | | / | |
| **fig6-d\e\f** | image patches from 2024111404-1-2 | | 20nm*20nm*10nm<br><br>1024*884 | |
| **fig6-h** | 2024111404-1-2 3D cell volume | | / | |
